# Targeting Pulmonary Fibrosis by SLC1A5 dependent Glutamine Transport Blockade

**DOI:** 10.1101/2022.05.23.493168

**Authors:** Malay Choudhury, Kyle J. Schaefbauer, Theodore J. Kottom, Eunhee S. Yi, Daniel J. Tschumperlin, Andrew H. Limper

## Abstract

The neutral amino acid glutamine plays a central role in TGF-β-induced myofibroblast activation and differentiation. Cells take up glutamine mainly through a transporter expressed on the cell surface known as solute carrier SLC1A5. In this current work, we demonstrated that profibrotic actions of TGF-β are mediated, at least in part, through a metabolic maladaptation of SLC1A5 and targeting SLC1A5 abrogates multiple facets of fibroblast activation. This approach could thus represent a novel therapeutic strategy to treat fibroproliferative diseases. We found that SLC1A5 was highly expressed in fibrotic lung fibroblasts and fibroblasts isolated from IPF lungs. The expression of profibrotic targets, cell migration, and anchorage independent growth by TGF-β required the activity of SLC1A5. Loss or inhibition of SLC1A5 function enhanced fibroblast susceptibility to autophagy, suppressed mTOR, HIF, Myc signaling, and impaired mitochondrial function, ATP production and glycolysis. Pharmacological inhibition of SLC1A5 by small molecule inhibitor V-9302 shifted fibroblast transcriptional profiles from profibrotic to fibrosis resolving, and attenuated fibrosis in a bleomycin treated mouse model of lung fibrosis. This is the first study, to our knowledge, to demonstrate the utility of a pharmacological inhibitor of glutamine transport in fibrosis, laying a framework for new paradigm-shifting therapies targeting cellular metabolism for this devastating disease.

## Introduction

Fibroproliferative diseases are a leading cause of morbidity and mortality featuring localized and systematic tissue/organ fibrosis (1). Nearly 45% of all deaths in the developed world are caused by chronic inflammatory and fibrogenic disorders (1). Idiopathic Pulmonary Fibrosis (IPF) is a debilitating, chronic, and irreversible Interstitial Lung Diseases (ILD) with unclear etiology, characterized by aberrant accumulation of extracellular matrix proteins (ECM) in the lungs of individuals between 50 and 85 years old, leading to respiratory failure and death typically 2-5 years after diagnosis (1–3). The incidence of IPF has doubled over the decade (4) and an estimated 50,000 people die each year from IPF in the U.S, more deaths than many cancers including breast cancer (5). Although two FDA approved drugs for IPF, Nintedanib and Pirfenidone (6), have been shown to slow the progression of disease, there are a significant number of side effects which limit therapeutic benefit over time (7). Thus, there is an urgent need to better understand the molecular mechanisms and pathophysiological processes driving IPF to identify novel therapeutic targets and discover effective treatment strategies to improve patient outcomes.

Proliferating cells face a challenge in balancing appropriate level of amino acids required for macromolecular synthesis and growth. As glutamine is the most abundant amino acid in plasma, renewed interest in glutamine metabolism during fibrosis was sparked by the recognition that glutaminolysis controls activation of myofibroblast in hepatic fibrosis, nonalcoholic steatohepatitis, cardiac fibrosis and iatrogenic laryngotracheal stenosis through upregulation of glutaminase 1(GLS1), Yes-associated protein 1 (YAP1) or nuclear factor-erythroid 2-related factor2 (Nrf2) (8–11). Moreover, recent reports including our work have shown that GLS1 which converts glutamine to glutamate plays a central role in TGF-β-induced myofibroblast activation and differentiation and in bleomycin-induced experimental pulmonary fibrosis (12–16). Although promising, recent studies also showed that pharmacological inhibition of GLS1 is ineffective in targeting tumors in vivo, even with elevated expression of GLS1 mainly due to metabolic adaptations which renders glutaminase dispensable (17, 18). As a hydrophilic amino acid, extracellular glutamine cannot penetrate the plasma membrane without the aid of cell surface glutamine transporters i.e., solute carrier proteins (SLCs) (19). ASCT2, encoded by solute carrier transporter 1A5 (SLC1A5) is a Na +-dependent bidirectional amino acid transporter that mainly mediates cellular uptake of large neutral amino acids specifically glutamine (20). Additional substrates transported by SLC1A5 include alanine, serine, cysteine, threonine, leucine, and asparagine (21). Several studies further demonstrated that SLC1A5 is required for the growth of a variety of cancers and targeting SLC1A5 by either RNAi or small molecule inhibitors suppresses cancer cell proliferation in vitro and tumor growth in vivo (21–29). The advantages of SLC1A5 as a drug target are: (i) SLC1A5 is expressed on the cell surface and are therefore targetable by both small molecules and therapeutic antibodies; (ii) given that SLC1A5 transport metabolites to the rapidly-growing cells, inhibiting SLC1A5 for even a short period of time, would starve fibrotic fibroblasts while sparing healthy cells; and (iii) amino acid transporters have deep binding pockets, which are ideal sites for the multiple binding of inhibitors. However, little is known currently about the biological impact and mechanism of regulations of SLC1A5 in pulmonary fibrosis.

In the present study, we have found that SLC1A5 was highly expressed in fibrotic lung fibroblasts and the expression of profibrotic targets, cell migration, and anchorage independent growth by TGF-β required the activity of SLC1A5. Loss or inhibition of SLC1A5 function suppresses mTOR, HIF, cMyc signaling, and impairs mitochondrial function, ATP production and glycolysis. Pharmacological inhibition of SLC1A5 shifts the profibrotic phenotype to a fibrosis resolving phenotype in the bleomycin-treated mouse model of lung fibrosis. Therefore, our results support a novel molecular and metabolic pathway modulating fibrosis resolution and represent a potential new paradigm-shifting approach for therapy targeting cellular metabolism.

## Results

### SLC1A5 expression is upregulated in pulmonary fibrosis

In this current research, we investigated the interrelationship between glutamine transporter(s) and profibrotic TGF-β signaling. Glutamine is transported into cells through plasma membrane glutamine transporters such as SLC1A5, SLC38A1 and SLC38A2 (29). In primary human lung fibroblasts cells (NHLF), qPCR analysis of transcripts for the important glutamine transporter(s) (SLCs) showed only SLC1A5 expression was dramatically elevated by TGF-β (Figure 1A), and this finding was confirmed and extended by western blot and qPCR analysis where elevated expression of SLC1A5 protein and mRNA was observed by TGF-β in NHLF in a time dependent manner (Figure 1B and 1C). This was specific to TGF-β stimulation as treatment with the TβRI kinase inhibitor, SB431542, abolished the response in NHLF (Figure 1D). In addition, fibroblasts isolated from lungs of IPF patients also showed significantly enhanced basal SLC1A5 levels compared to primary human lung fibroblasts (Figure 1E and 1F). We further investigated whether SLC1A5 knockdown upregulated other amino acid transporter(s). We did not observe any significant upregulation by TGF-β on other glutamine transporters upon SLC1A5 knock down confirming that SLC1A5 suppression is without significant transporter plasticity or redundancy (Figure 1G). In addition, enhanced SLC1A5 expression was observed in mice that were subjected to a fibrosis-inducing bleomycin treatment (Figure 1H).

**Figure 1.**
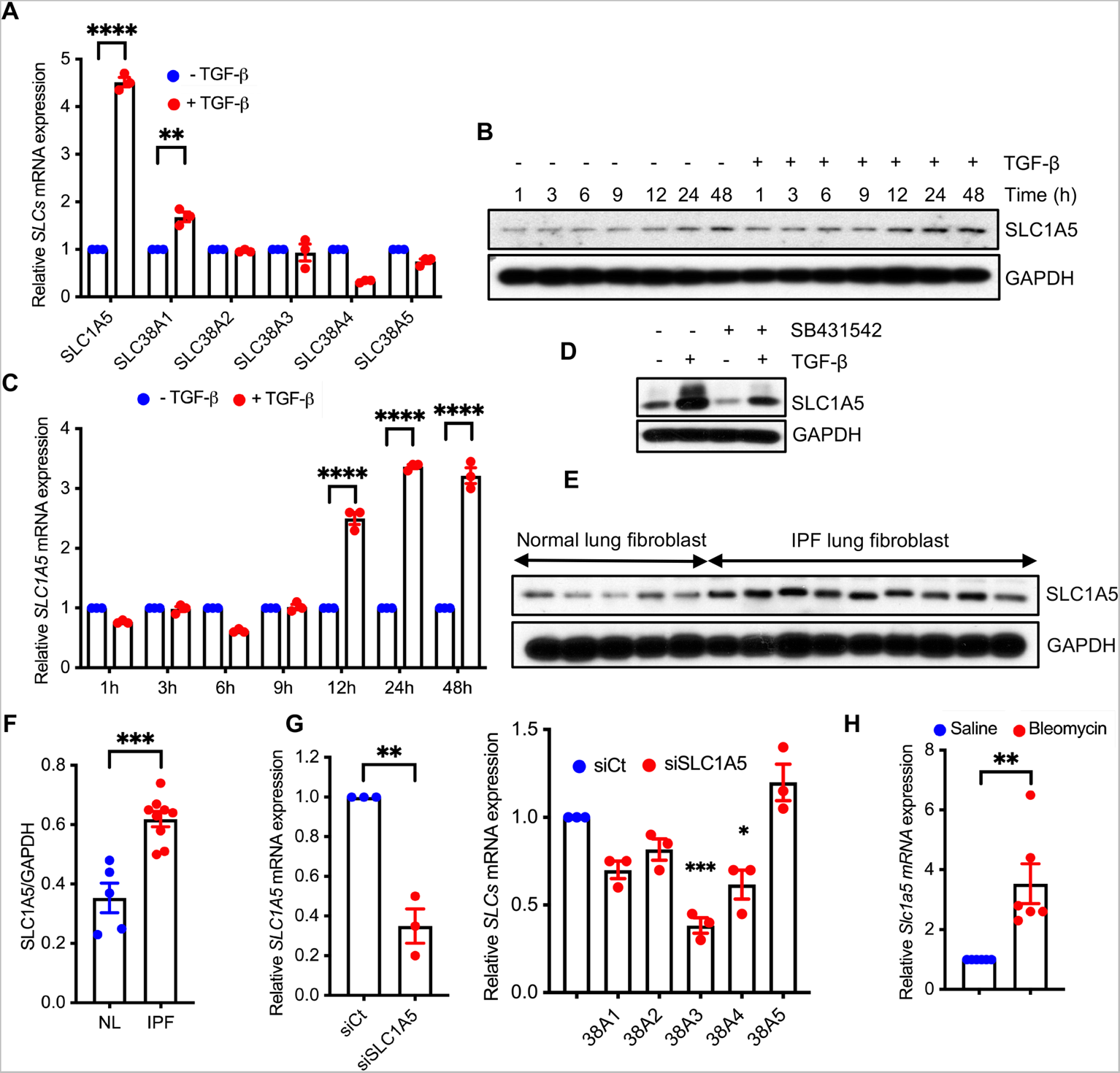
TGF-β stimulates SLC1A5 and its expression upregulated in fibrosis. (**A**) Normal Human Lung Fibroblasts (NHLF) cells were treated with 5 ng/ml TGF-β or vehicle (4 mM HCL + 10 mg/ml BSA) for 48h and qPCR analysis of above *SLCs* were performed. (**B**) Western blot analysis of SLC1A5 was determined in quiescent NHLF cells in the absence (−) or presence (+) of 5 ng/ml TGF-β at the indicated times. (**C**) NHLF cells were treated as in (B) and 1μg RNA was subjected to cDNA synthesis and subsequently qPCR analysis using *SLC1A5* primers. (**D**) NHLF cells were treated with DMSO (0.1%) or the TbRI inhibitor SB431542 (10 μM) with or without TGF-β (5 ng/ml, 48h induction) and protein expression of SLC1A5 was determined by Western blotting. (**E**) Primary lung fibroblasts from patients with IPF and healthy controls were propagated for 3-4 passages in cell culture and Western blot analysis for SLC1A5 was performed. (**F**) Ratios of SLC1A5 to GAPDH in normal and IPF lung fibroblasts. (**G**) qRT-PCR analysis of important amino acid transporter genes in SLC1A5 knocked down NHLF cells. (**H**) Basal SLC1A5 expression from lungs of saline and bleomycin treated mice by qPCR analysis (n = 6). Data generated for qRT-PCR represent mean ± SEM of n = 3 independent experiments. Western blots are representative of 3 independent experiments. Differences between groups were evaluated by two-way (A, C) or one-way (G) ANOVA test with Tukey post-hoc analysis or unpaired two-tailed student’s t-test (F, H) using GraphPad Prism 9.3 software. **P < 0.01, ***P < 0.001, ****P < 0.0001.

### Profibrotic TGF-β signaling is dependent on SLC1A5

We next investigated the role of SLC1A5 in profibrotic TGF-β signaling. SLC1A5 expression using siRNA or SLC1A5 activity using V-9302 or GPNA was inhibited and TGF-β induction of profibrotic molecules was determined by western blotting and qPCR in NHLF (Figure 2A-2E), IPF fibroblast (Figure 2F) and AKR-2B (a nontransformed murine fibroblast) (Supplementary Figure 1A-1D) cells. V-9302 is a newly developed selective inhibitor of SLC1A5 which was effective in suppressing tumor proliferation via inhibition of mTOR signaling (21). GPNA (L-γ-Glutamyl-p-nitroanilide) is a potent inhibitor of SLC1A5 as shown previously (22, 30). We performed MTT and XTT assay (cell viability and proliferation assay) which confirmed no significant effect of V-9302 on cellular proliferation (i.e., toxicity) at 10μM concentrations that significantly impacted profibrotic TGF-β signaling (Supplementary Figure 2A and 2B). This concentration was subsequently used throughout this study as reported in different tumor models (28, 31, 32). There were also no significant changes in gene expression predominantly associated with DNA repair (*ATM, ATR* and *BRCA1*), recombination (*RAD51* and *RAD52*) and cell cycle (*CCNB1* and *CCND1*), under V-9302 treatment conditions (Supplementary Figure 2C). While phosphorylation of SMAD2 and SMAD3 occurs independently of SLC1A5 (Supplementary Figure 3), expression of profibrotic targets including Col1(Collagen I), FN (Fibronectin), CTGF (Connective tissue growth factor) and ACTA2 (alpha smooth muscle actin) requires the action SLC1A5 (Figure 2A-2F and Supplementary Figure 1A-1D) only in fibroblasts as we did not see any detectable expression of profibrotic markers in human epithelial or murine alveolar macrophage cells by TGF-β treatment (Supplementary Figure 4A and 4B). Next, we tested whether exogenous cell permeable a-KG which is downstream of SLC1A5 in glutamine metabolism pathway, can restore TGF-β mediated induction of profibrotic markers. The addition of a-KG overcame the inhibitory effect of SLC1A5 knockdown on the induction of Col1, FN, and ACTA2 by TGF-β (Figure 2G). Collagen facilitates cell–ECM interactions and undergoes crosslinking in the extracellular space. Lysyl oxidase (LOX) primarily mediates this process and form cross links in extracellular matrix proteins. LOX expression is increased in multiple fibrotic diseases and accompanied by increased stiffness and decreased ECM degradation (33). We investigated whether V-9302 treatment altered the expression of key matrix cross-linking and degradation genes. V-9302 treatment downregulated mRNA expression of key matrix cross-linking genes (Figure 2H) whereas expression of key genes associated with matrix degradation and clearance in IPF fibroblasts were upregulated (Figure 2I). Together, these data suggest that myofibroblast differentiation in part depends on the availability of SLC1A5 in TGF-β mediated lung fibrosis.

**Figure 2.**
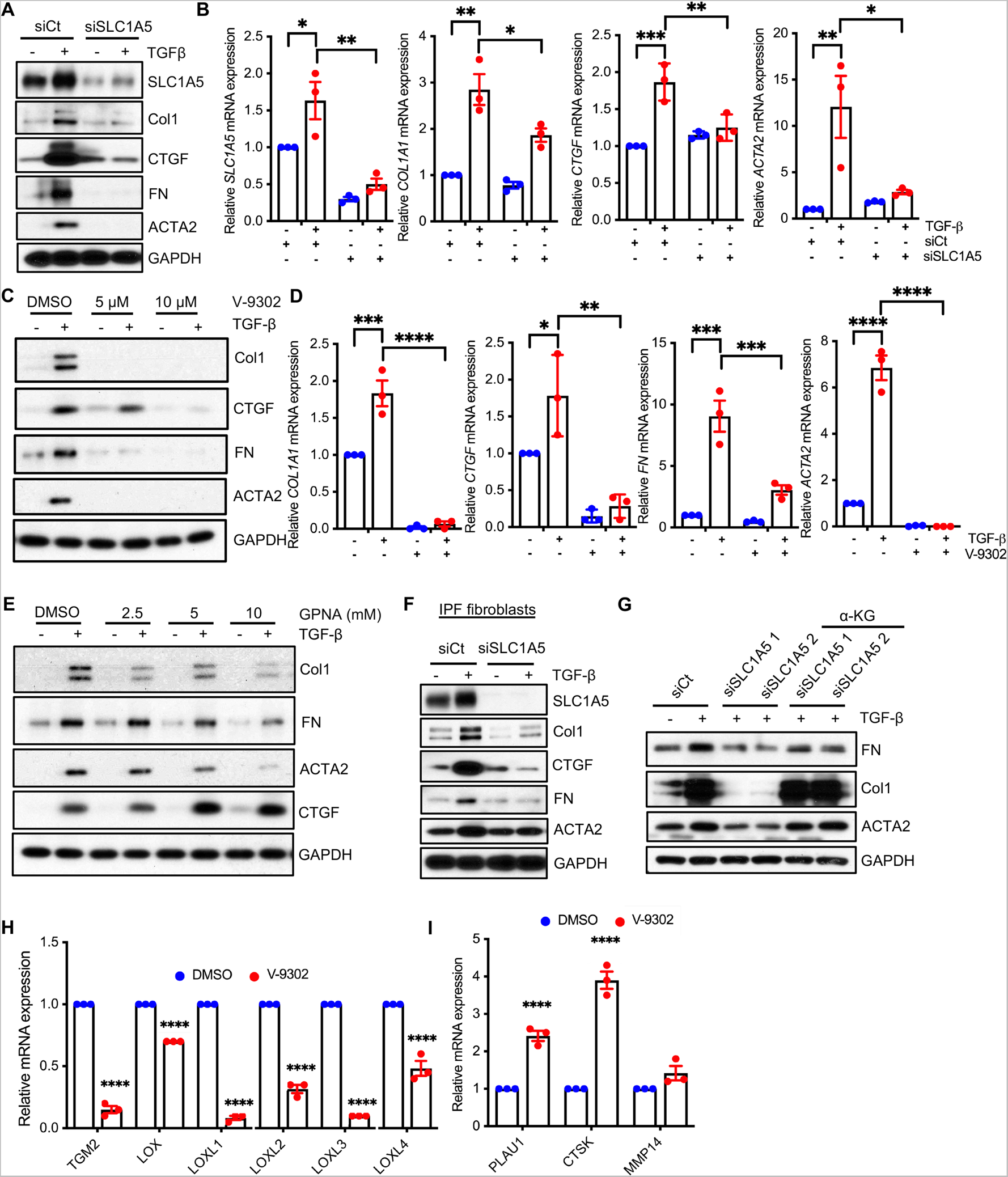
Profibrotic TGF-β signaling is dependent on the action of SLC1A5. (**A, B**) NHLF cells were transfected with non-targeting control (siCt) or siRNA against SLC1A5 (siSLC1A5). Vehicle (−) or TGF-β (+) was directly added to a final concentration of 5 ng/ml and following 48h incubation, Western blotting (A) and qPCR (B) of profibrotic molecules were performed. (**C**) Quiescent NHLF cells in 0.1% FBS/DMEM were pretreated for 1h with Vehicle (0.1% DMSO) or 5 and 10 μM V-9302 (SLC1A5 inhibitor) prior to addition of Vehicle (-; 4 mM HCL + 10 mg/ml BSA) or TGF-β (+; 5 ng/ml). Following 48h incubation, lysates were prepared and Western blotted for Col1, CTGF, FN and ACTA2. GAPDH was used as a loading control. (**D**) qPCR of the indicated genes was performed as in (C) (10 μM V-9302). (**E**) Quiescent NHLF cells treated with 0.1% DMSO or 2.5, 5 and 10 mM GPNA for 1h with the presence or absence of TGF-β (5 ng/ml) for another 48h. After 48h, cell lysates were prepared and Western blotted for indicated proteins. (**F**) IPF fibroblasts were transfected with either non-targeting control (siCt) or siRNA against SLC1A5 (siSLC1A5) and Western blotted as mentioned in Fig. 2A. (**G**) NHLF cells were transfected with control or SLC1A5 siRNA, treated with vehicle or diethyl-2-oxopentanedioate (esterified form of α-KG, 2 mM) for 12h, and then stimulated with vehicle or TGF-β for 24h before Western blotting for the indicated proteins. (**H, I**) NHLF were treated with Vehicle (0.1% DMSO) or 10 μM V-9302 for 48h. Western blotting was performed for ECM crosslinking genes (H) and ECM degradation genes (I). qRT-PCR represent mean ± SEM of n = 3 independent experiments. The Western blots are representative of 3 independent experiments. *P < 0.05, **P < 0.01, ***P < 0.001, ****P < 0.0001 were calculated by two-way ANOVA test with multiple comparison by Tukey post-hoc analysis using GraphPad Prism 9.3 software.

### Glutamine uptake, Cell migration and Anchorage Independent Growth (AIG) are dependent on SLC1A5 activity

As SLC1A5 is a high affinity glutamine transporter, we sought to investigate the metabolic impact of SLC1A5 inhibition in vitro. To examine the inhibition of glutamine transport by V-9302 in NHLF, we measured Intracellular glutamine level. We found a marked reduction of intracellular glutamine uptake in SLC1A5 inhibited cells (Figure 3A). The preceding findings were further extended to examine the effect of SLC1A5 inhibition on cell migration and anchorage independent growth (AIG) stimulated by TGF-β. SLC1A5 knockdown or inhibition in NHLF and AKR-2B cells significantly reduced TGF-β stimulated cell migration (Figure 3B, 3C and Supplementary Figure 5). A similar requirement for SLC1A5 activity was observed for TGF-β stimulated anchorage independent growth in soft agar (Figure 3D).

**Figure 3.**
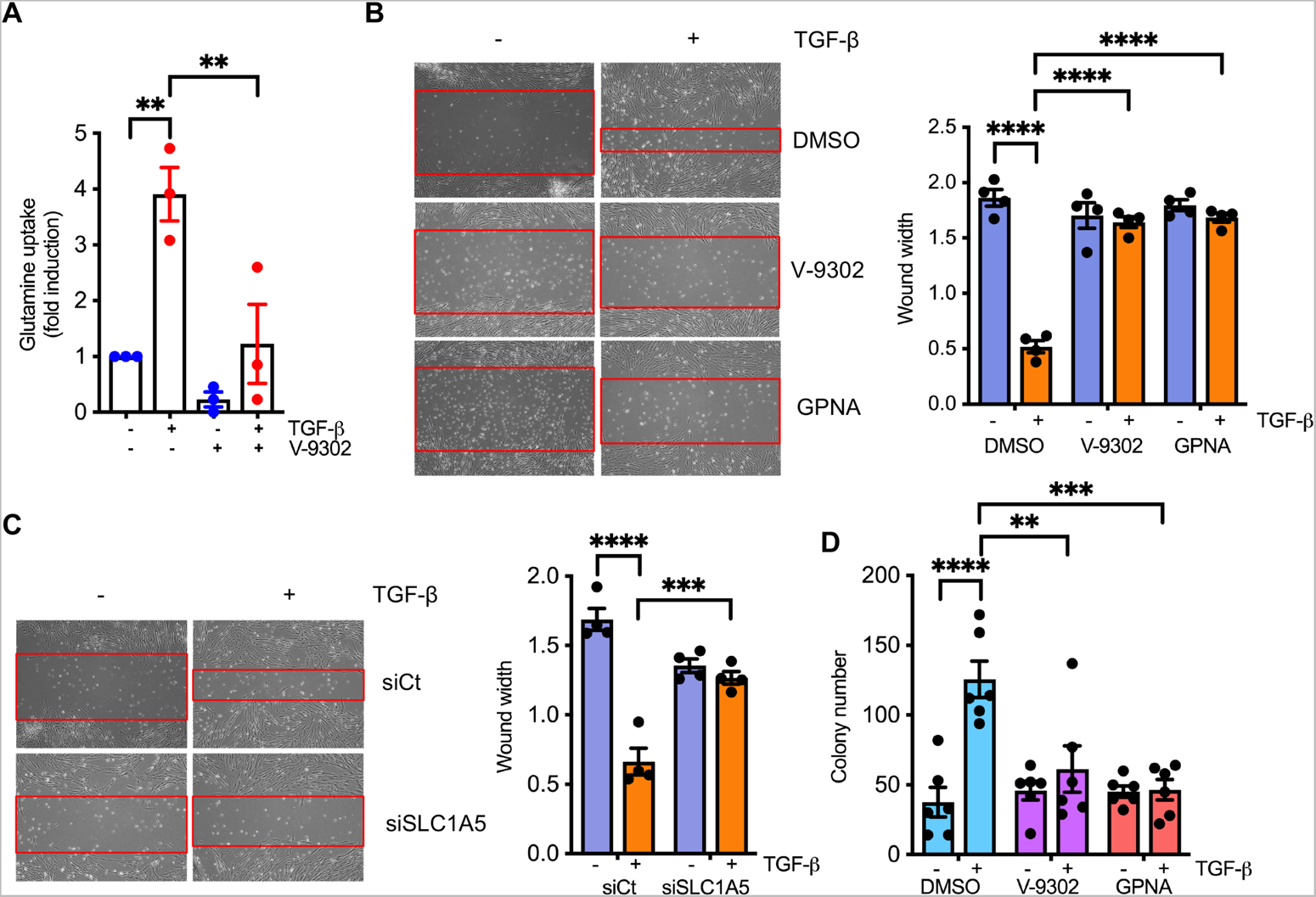
Glutamine uptake, Cell migration and Anchorage Independent Growth (AIG) are dependent on SLC1A5 activity. (**A**) Glutamine uptake assay in TGF-β –treated (24h) NHLF cells. Results show an increase in glutamine uptake by TGF-β, which is inhibited by the V-9302 (10µM). (**B**) (Left) Scratch assays were performed on NHLF cells. Red bands indicate the leading edge following 24h in the presence (+) or absence (−) of TGF-β (5 ng/ml) alone or containing V-9302 (10 μM) or GPNA (10mM) and are representative of 3 separate experiments. (Right) Quantification of wound closure. Data reflect mean ± SEM of n = 3. (**C**) (Left) NHLF cells were transfected with control or SLC1A5 siRNA and then subjected to scratch assay as in (B). Red lines indicate the leading edge after 24h and representative of 3 separate experiments. Data represents mean ± SEM of n = 3. (Right) Quantification of wound closure. (**D**) Inhibition of SLC1A5 activity by V-9302 and GPNA significantly affects anchorage-independent growth of NHLF cells. Quantitative analyses of cell colonies in soft agar are shown. Data reflects mean ± SEM of n = 6. **P < 0.01, ***P < 0.001, ****P < 0.0001 were calculated by two-way ANOVA test with Tukey post-hoc test using GraphPad Prism 9.3 software.

### SLC1A5 deficiency suppresses mTOR signaling and leads to transcriptional reprogramming towards survival

SLC1A5 was previously shown to be essential for mTOR activation (21). We explored whether SLC1A5 loss caused a reduction in the phosphorylation of mTOR targets. Consistent with this hypothesis, silencing of SLC1A5 or inhibition by V-9302 or GPNA resulted in markedly decreased p-S6K, p-Akt, p-4E-BP1 levels, suggesting glutamine deprivation inhibits the mTOR pathway in fibrosis (Figure 4A, 4B and Supplementary Figure 6). Although TGF-β signals mainly via the SMAD pathway, it also activates other pathways collectively referred to as ‘non-canonical’ signaling (PI3K/AKT/mTOR) which often complement SMAD action (34–37). The canonical SMAD signaling pathway is activated within minutes of TGF-β addition, whereas activation of non-SMAD pathway requires much longer treatment times (15, 35, 38). As SLC1A5 induction occurs relatively late (Figure 1B and 1C), it would not be surprising for TGF-β to stimulate SLC1A5 through both canonical and noncanonical mechanisms. To further identify the pathway(s) required for the induction of SLC1A5 by TGF-β, NHLF cells were treated, first, with siRNA that targeted SMAD2, SMAD3, or a non-targeting control, (Figure 4C); or secondly, with MAPK kinase (MEK), PI3K, AKT, mTORC1 and mTORC1/2 inhibitors, U0123, LY294002, MK2206, rapamycin and Torin 1 respectively (Figure 4D) and the induction of SLC1A5 by TGF-β was measured. SMAD2 or, SMAD3 knockdown as well as Inhibition of PI3K and mTORC1/2 abrogated the induction of SLC1A5 by TGF-β (Figure 4C and 4D). As activation of autophagy is known to provide crucial source of nutrient in response to nutrient depletion, SLC1A5 deficient NHLF cells showed a significant induction of autophagy as demonstrated by elevated expression of microtubule-associated protein 1 light chain 3B II (LC3BII) and beclin1 (Figure 4E-4G). We further confirmed that endoplasmic reticulum stress and unfolded protein response genes were also activated as a compensatory response following inhibition of SLC1A5 (Figure 4H).

**Figure 4.**
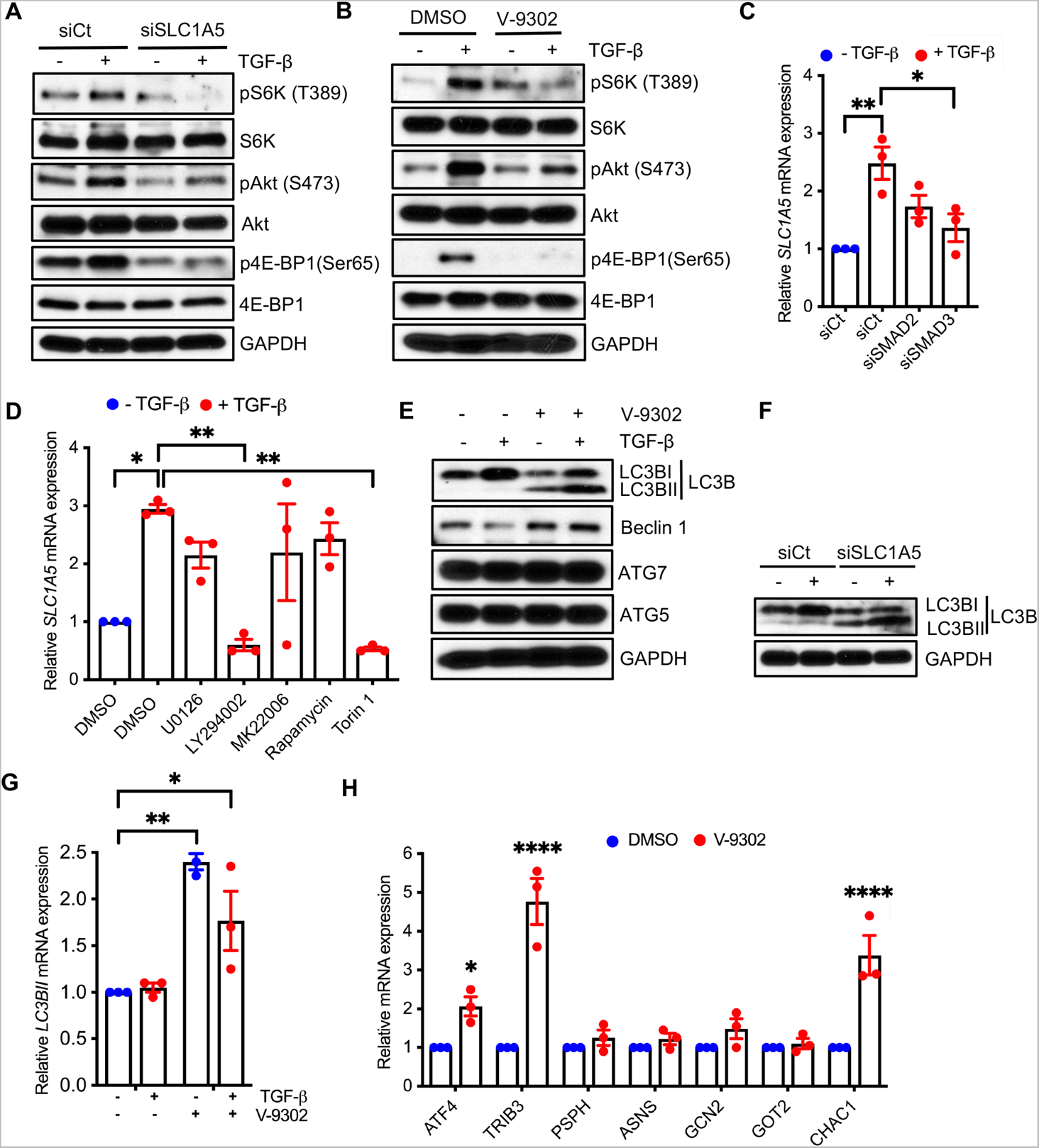
SLC1A5 deficiency suppresses mTOR signaling and leads to transcriptional reprogramming towards survival. (**A**) NHLF cells were transfected with scrambled control or SLC1A5 siRNA and Western blotted for the indicative proteins 6h post TGF-β or vehicle treatment. (**B**) NHLF cells were treated with V-9302 and stimulated in the absence (−) or presence (+) of TGF-β (5 ng/ml) and indicated proteins were assessed at 6h. (**C**) NHLF cells with either non-targeting control (siCt) or siRNA targeting SMAD2 or SMAD3 were treated with TGF-β (+; 5 ng/ml) or Vehicle (−) and qPCR of *SLC1A5* was performed 24h post treatment. (**D**) qPCR for *SLC1A5* 24h post Vehicle (−) or TGF-β (+) treatment in the presence of 0.1% DMSO; MEK-ERK1/2 inhibitor U0126 (3 μM); PI3K inhibitor, LY294002 (20 μM); Akt inhibitor, MK22006 (300 nM); mTORC1 inhibitor, Rapamycin (100 nM); or mTORC1+C2 inhibitor, Torin 1 (200 nM). (**E, F, G**) NHLF cells were treated as of Fig. 2A and 2C and Western blot analysis of autophagic proteins (E, F) or qPCR of *LC3BII* (G) were performed. (**H**) qRT-PCR analysis of endoplasmic reticulum stress/unfolded protein response genes in SLC1A5 inhibited NHLF cells. All Western blots are representative of 3 separate experiments. Data in (C, D, G and H) represent mean ± SEM and n = 3 independent experiments. *P < 0.05, **P < 0.01, ****P < 0.0001 were calculated by one-way (C, D) or two-way (G, H) ANOVA with Tukey post-hoc test using GraphPad Prism 9.3 software.

### Role of SLC1A5 in HIF and c-Myc signaling

Since hypoxia Inducing Factors (HIF) and cMyc controls the progression and pathogenesis of fibrosis (39–42) and cMyc upregulation has been associated with increased glycolysis and glutaminolysis to support the increased biosynthetic demands (43), we next determined whether these transcription factors participate in TGF-β–induced SLC1A5 expression. Silencing HIF1α, HIF2α and cMyc expression impaired the protein expression of SLC1A5 (Figure 5A), whereas knock down or inhibition of SLC1A5 downregulated protein expression of HIF1α, HIF2α, and cMyc (Figure 5B and 5C). Furthermore, downregulation of SLC1A5 impaired the gene expression of HIF1α, HIF2α and cMyc (Figure 5D). Together the data support a mechanism by which HIF, cMyc and SLC1A5 constitute a positive feedback loop that sustain each other’s expression to control mTOR signaling and fibrosis development. These observations provide further rationale for targeting SLC1A5 as a potential therapeutic strategy for pulmonary fibrosis.

**Figure 5.**
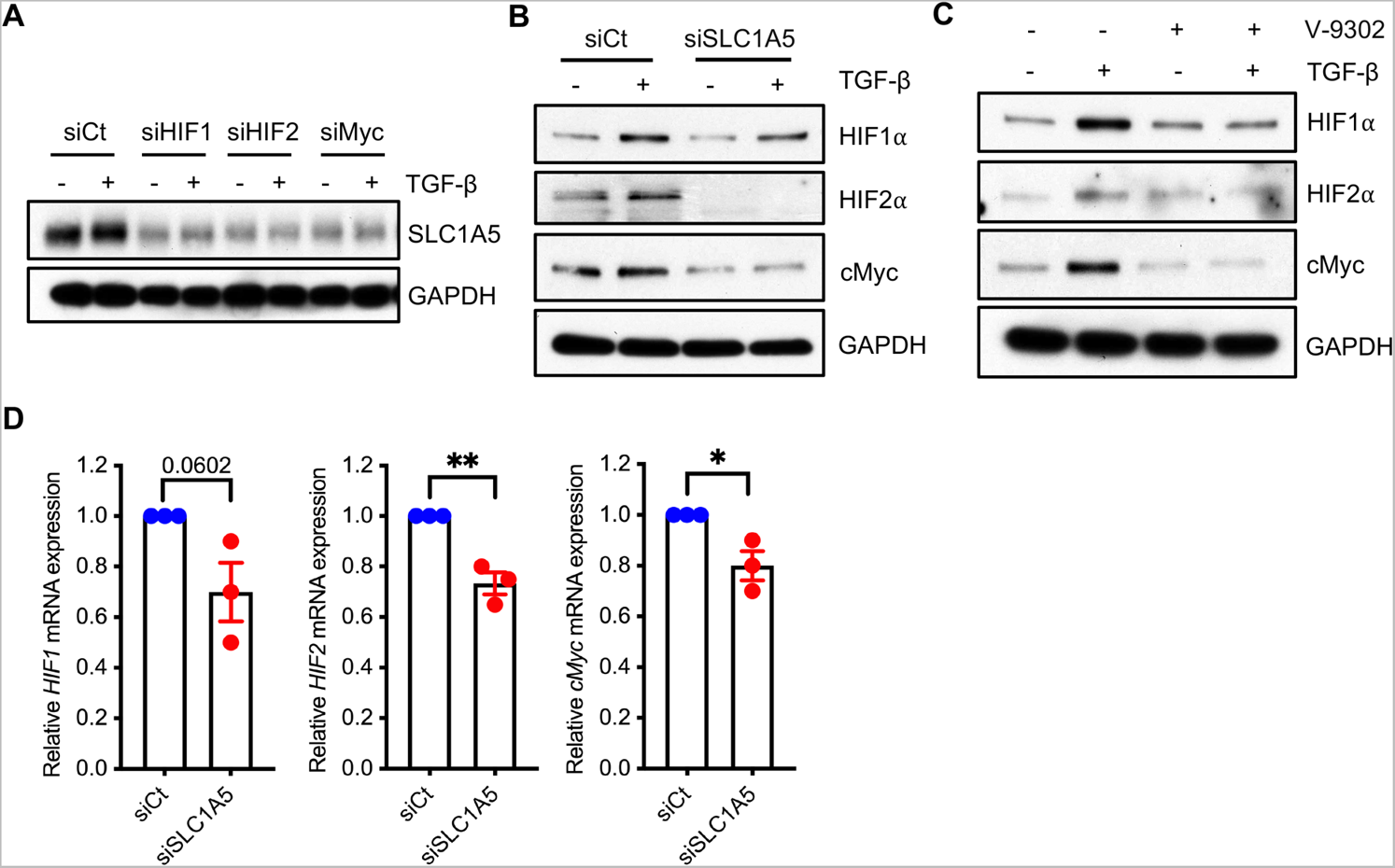
Role of SLC1A5 in HIF and c-Myc signaling. (**A, B**) NHLF cells were transfected with control or HIF1α, HIF2α and cMyc siRNA (A) or SLC1A5 siRNA (B). Following 48h incubation (+/− 5 ng/ml TGF-β), lysates were prepared and western blotted for indicated proteins. (**C**) NHLF cells were treated as of Fig. 2C, and Western blot analysis was performed. (**D**) qPCR analysis of *HIF1*, *HIF2* and *cMyc* genes in *SLC1A5* knocked down NHLF cells. Western blots are representative of 3 independent experiments. Differences between two groups was evaluated by unpaired two-tailed student’s t-test using GraphPad Prism 9.3 software. *P < 0.05, **P < 0.01.

### SLC1A5 is critical for metabolic reprogramming in fibroblasts

Myofibroblast differentiation requires metabolic reprogramming characterized by an increase in OXPHOS and glycolysis (44). To determine whether this metabolic reprogramming requires SLC1A5, we analyzed the inhibition of SLC1A5 following TGF-β induced differentiation on the oxygen consumption rate (OCR) and the extracellular acidification rate (ECAR) of myofibroblasts. Inhibition of SLC1A5 by V-9302 impaired the basal and maximal OCR, implying defective mitochondrial respiration (Figure 6A and 6B). Inhibition of SLC1A5 suppressed the ECAR (Figure 6C and 6D). Furthermore, rotenone and antimycin (mitochondrial complex I and III inhibitors respectively) treatment suppressed SLC1A5 and fibrosis marker expression, whereas 3-NP (mitochondrial complex II inhibitor) did not have a significant effect (Figure 6E). We also examined the effect of SLC1A5 expression on mitochondria derived ATP production. Downregulation of SLC1A5 significantly decreased ATP production indicating ATP production is highly dependent on SLC1A5 (Figure 6F). TFAM (Mitochondrial transcription factor A) is involved in mitochondrial DNA transcription and replication and silencing of TFAM down-regulates TGF-β-induced expression of ACTA2 (44). As TFAM is important in maintaining mitochondrial fitness, we examined the roles of mitochondrial fitness and metabolism by silencing TFAM and determined the effect on the expression of SLC1A5. TFAM silencing decreased expression of SLC1A5 (Figure 6G). Silencing of SLC1A5 markedly reduced the expression of TFAM expression in NHLF (Figure 6H). Taken together, these data suggests that both mitochondrial biogenesis and SLC1A5 expression coordinately support myofibroblast differentiation.

**Figure 6.**
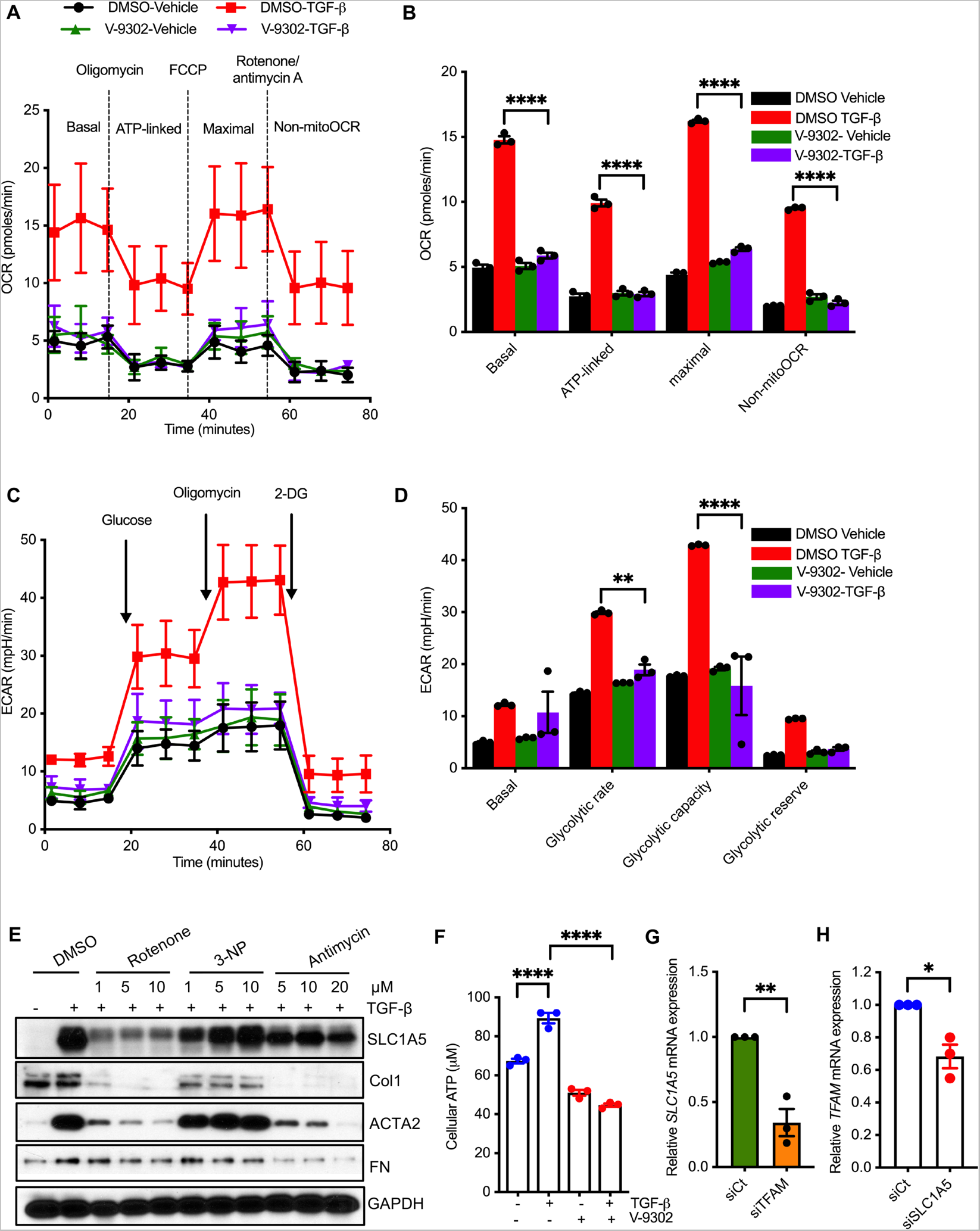
SLC1A5 is critical for metabolic reprogramming in fibroblasts. (**A, B**) Quiescent NHLF cells were treated for 24h with vehicle (DMSO) or V-9302 in the presence or absence of TGF-β, and oxygen consumption rate (OCR) was assessed by the Seahorse XFp Cell Mito Stress Test Kit on a Seahorse XFp extracellular flux analyzer. Data are presented as individual time points (A) and as averages (B). (**C, D**) NHLF cells were treated as in (A) and then glycolysis was measured as ECAR by Seahorse XF Glycolysis Stress Test Kit. Data represents individual time points (A) and averages (B) (**E**) NHLF cells were pretreated for 1h with Vehicle (0.1% DMSO) or rotenone, 3-NP and antimycin (mitochondrial complex I, II and III inhibitors respectively) prior to addition of Vehicle (-;) or TGF-β (+; 5 ng/ml). Following 48h incubation, lysates were prepared and Western blotted for SLC1A5 and fibrosis marker expression. (**F**) NHLF cells were treated as of Fig. 2C, and cellular ATP level were measured. (**G, H**) NHLF cells were transfected with NT or TFAM siRNA (G) or SLC1A5 siRNA (H) and expression levels of *SLC1A5* and *TFAM* were measured. n = 3 independent experiments. All data reflect the means ± SEM. Differences between groups were evaluated by two-way ANOVA with Tukey post hoc analysis (B, D, and F) or unpaired Student’s t test (G, H). *P < .05, **P < 0.01, ****P < 0.0001.

### V-9302 treatment in a therapeutic murine model ameliorates bleomycin induced fibrosis

In that SLC1A5 has been successfully targeted by V-9302 in several tumor models with an optimal dose of 50-75 mg/kg/day (21, 28), we extended our findings to a murine treatment model of lung fibrosis. C57BL/6 male and female mice were intratracheally treated with an equal volume of saline (control) or bleomycin (BLM) (3.5 U/Kg). On day 11, BLM and saline treated mice started receiving treatment of Vehicle (PBS supplemented with 2% DMSO) or V-9302 (37.5 mg/kg/day) daily by intraperitoneal injection for 13 days and then euthanized the animals on day 25 (Figure 7A). Peripheral blood oxygen saturation on room air (SpO2) using pulse oximeter provides a useful non-invasive measure of overall lung function and morbidity or mortality for experimental pulmonary fibrosis. To accurately measure lung function throughout the study, SpO2 was determined before and every third or fourth day after the initiation of treatment on day 11. V-9302 stabilized or improved peripheral blood oxygenation during treatment (Figure 7B). Flexivent analysis of lung compliance was determined following euthanasia on day 25. FlexiVent analysis indicated a V-9302 dependent improvement of lung compliance (Figure 7C). These physiologic findings further supported beneficial effects of V-9302 (i) lung weight at study completion (Figure 7D); (ii) reduced collagen deposition in the lung, as assessed by both hydroxyproline content (Figure 7E); (iii) H&E and trichrome staining by V-9302 treatment (Figure 7F and 7G and Supplementary Figure 7); (iv) reduced expression of profibrotic markers following V-9302 treatment (Figure 7H). Importantly, V-9302 treated mice did not show any body weight difference compared to control mice (21, 45). V-9302 also had no demonstrable effect on murine liver and kidney function (Supplementary Figure 8 and 9) or blood count parameters (Supplementary Figure 10). Taken together, these results demonstrate that pharmacological inhibition of Slc1a5 with V-9302 during the fibrotic phase is sufficient to attenuate bleomycin induced pulmonary fibrosis. Since sex may impact fibrosis development and resolution, development of fibrosis and their resolution was analyzed **in** both male and female mice to detect any potential differences. In our data, we observed that the effects of V-9302 **in** regulating fibrosis are independent of sex.

**Figure 7.**
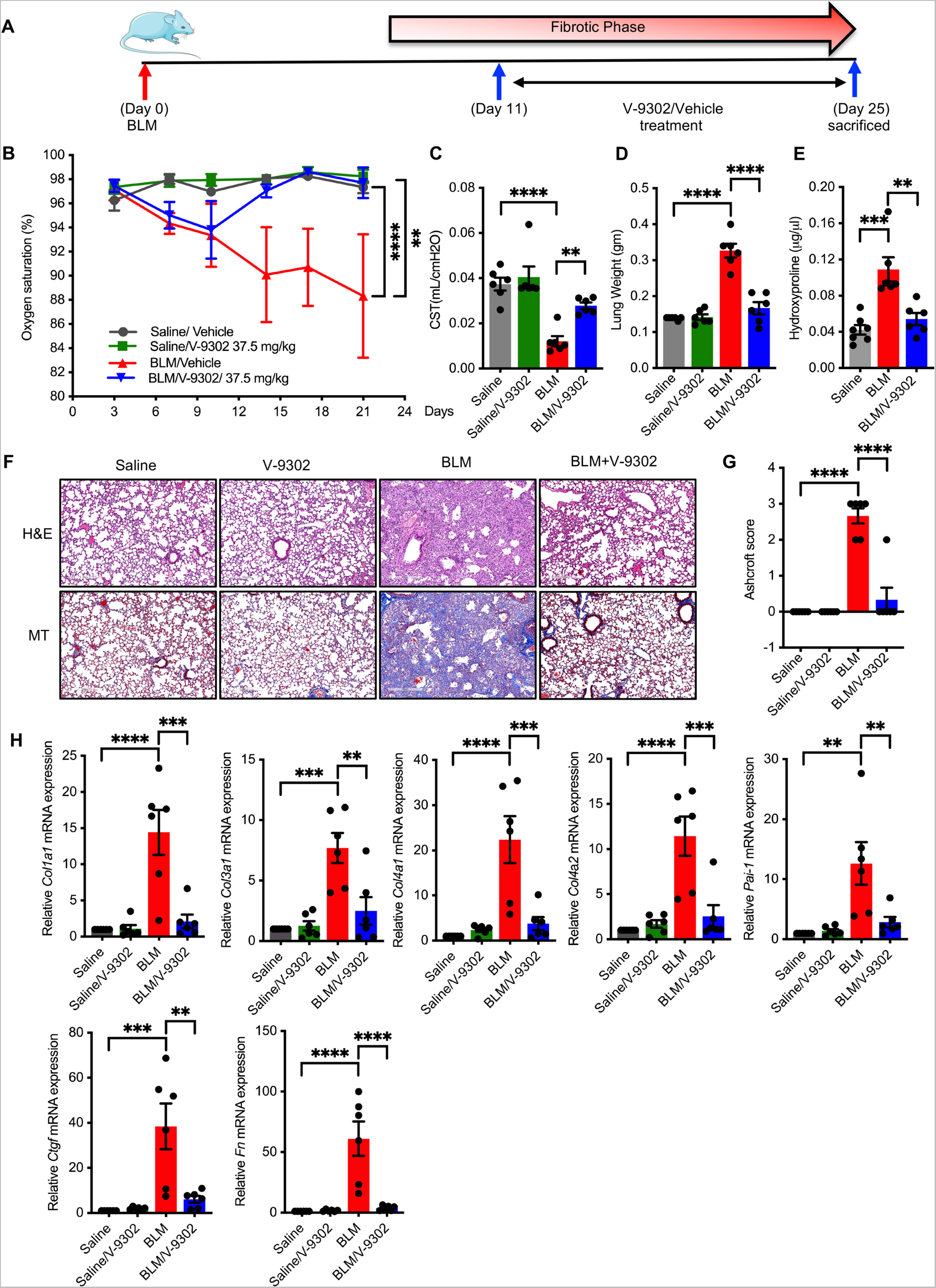
V-9302 treatment in a therapeutic murine model ameliorates bleomycin induced fibrosis. (**A**) Timelines for in vivo administration of bleomycin and V-9302. C57BL/6 male or female mice were intratracheally treated with an equal volume of saline (Control) or BLM (3.5 U/Kg). From day 11 to 24 mice were treated daily with either vehicle (PBS supplemented with 2% DMSO) or V-9302 (37.5 mg/Kg) by IP injection. On day 25, mice were euthanized. (**B**) Oxygen saturation levels was determined on every 3^rd^ day after BLM administration. (**C, D**) On day 25, mice were euthanized, and lungs subjected to Flexivent determination (C) and Total lung weight (D). (**E**) Total collagen content determined by hydroxyproline assay. (**F, G)** Hematoxylin and Eosin (H&E) staining for histology and Masson’s trichrome (MT) for collagen deposition (blue) were performed (F) and the lung histopathology sections were blindly scored for the degree of fibrosis using Ashcroft method (G). Scoring was performed based on the following criteria: 0 (no fibrosis), 1 (focal/minimal fibrosis), 2 (multifocal/moderate fibrosis), 3 (confluent/severe fibrosis), Scale bars, 300 μm. (**H**) qPCR analysis for fibrotic markers, *Col1a1, Col3a1, Col4a1*, *Col4a2, Pai-1, Ctgf,* and *Fn* in murine lung tissue harvested on day 25. Data reflect means ±SEM of 6 mice for each group (3 male and 3 female). Differences between groups were determined by two-way ANOVA test with Tukey post-hoc analysis (B) or one-way ANOVA with Tukey post-hoc analysis (C, D, E, G, H, I and J) using GraphPad Prism 9.3 software. **P < 0.01, ***P < 0.001, ****P < 0.0001.

## Discussion

Metabolic dysregulation contributes to the development of chronic lung diseases, including pulmonary fibrosis and its reliance on glutamine for energy supply is now well acknowledged. Recent reports by our group and others have shown that mitochondrial glutaminase, GLS1 plays a central role in TGF-β induced myofibroblast activation and differentiation in pulmonary fibrosis (12, 13, 15, 16). CB-839 (Calithera Biosciences) is a selective GLS1 inhibitor now being explored in Phase II clinical trials in multiple liquid and solid tumors (46). Although promising, limitations of this strategy include; (i) the presence of more than one isoform of GLS; (ii) unsolved subcellular localization that can hamper drug availability; (iii) GLS1 inhibition does not address the extra-mitochondrial roles of glutamine including MAPK signaling, the activity of GLS2 and amino acid transporters that require glutamine antiport for their function (for example SLC7A5) (21, 47–50). Therefore, blocking cellular glutamine transport would provide greater impact on glutamine metabolism rather than targeting downstream enzymes like GLS1 which provide extensive biological plasticity and redundancy to proliferating cells to maintain intracellular glutamine pool (21). Amino acid transporters such as SLC1A5 has been identified as the most critical plasma membrane glutamine transporter in cancer and is indispensable for active cell proliferation, mTOR signaling and Redox balance to name a few (51). Although a great deal of biological data documenting the medical importance of glutamine transport in cancer are known, there are no reports addressing how metabolic dysregulation by glutamine transporter(s) and cytokine autocrine/paracrine signaling function in an integrated and synergistic manner in the development of fibroproliferative diseases. Here we addressed these issues by (i) determining the role of SLC1A5 in profibrotic TGF-β signaling; (ii) investigating the mechanism(s) of cell type-specific induction of SLC1A5 by TGF-β and associated cellular, molecular, and metabolic mechanisms; (iii) evaluating how inhibition of SLC1A5’s action(s) “chemosensitizes” mesenchymal cultures to metabolic reprogramming; and, most importantly, (iv) extending these mechanistic findings to a treatment model of pulmonary fibrosis. To our knowledge, this is the first time the biological significance of SLC1A5 have been evaluated in fibrosis and indicate a potential therapeutic benefits of targeting SLC1A5.

Our data indicate an important relationship between the fibroproliferative actions of TGF-β and the induction of glutamine transporter, SLC1A5 in a cell type specific manner in fibroblasts. We extended these findings to include primary IPF fibroblasts to confirm the generality of the findings. Recently, Schulte et al., developed V-9302, a competitive small molecule antagonist that selectively and potently targets SLC1A5 (21). They further showed a stable V-9302–SLC1A5 interaction by drug affinity responsive target stability (DARTS) technique (52) and therefore implied that SLC1A5 is a V-9302 target. Pharmacological blockade of SLC1A5 with V-9302 resulted in attenuated cancer cell growth and proliferation, increased cell death, and increased oxidative stress, which collectively contributed to antitumor responses in vitro and in vivo (21). Herein, we showed that V-9302 inhibited SLC1A5-mediated glutamine uptake and using V-9302 or GPNA or siRNA mediated knock down of SLC1A5 inhibits fibrosis activation in primary human lung fibroblasts. As depletion of SLC1A5 significantly inhibits the import of essential amino acids (EAA) into cell, which might induce EAA shortage and trigger the amino acid stress response signal transduction pathway, we determined that SLC1A5 depletion increased ATF4, TRIB3 and CHAC1 expression (all involved in stress response pathway). An earlier report showed SLC1A5 deficiency was compensated by increased level of other glutamine transporter(s) SLC38A1/SLC38A2 in a GCN2 EIF2α kinase (GCN2) dependent, cell type specific manner (53).

Strikingly, we did not observe any similar increase of SLC38A1/SLC38A2 upon SLC1A5 knockdown confirming SLC1A5 is the main transporter of glutamine without significant transporter plasticity or redundancy in fibroblasts. It has been reported that activated fibroblasts self-amplify the fibrotic response by increasing matrix stiffness (54). We asked whether SLC1A5 inhibition influenced the expression of key matrix cross-linking and degradation genes. SLC1A5 antagonism resulted in a shift in transcriptional programs away from a contractile, proliferative, and matrix-depositing state toward a matrix-degrading and softening state potentially linked to fibrosis resolution.

We identified several molecular mechanism(s) associated with SLC1A5 knock down or V-9302 treatment. SLC1A5 down regulation or V-9302 exposure resulted in decreased mTOR activity as assessed by pS6K, pAkt, and p4EBP1, levels which is consistent with decreased amino acid metabolism and transport (21, 47). We further determined the role(s) of MAPK, PI3K, Akt, mTORC1, mTORC2, SMAD proteins (i.e., pathways all shown to be involved in profibrotic TGF-β signaling; (34, 55–59) in glutamine transport using a combination of genetic (siRNA) as well as pharmacologic approaches. Our data indicated that TGF-β signaling downstream of PI3K/mTORC2/SMAD2/3 are critical for the induction of SLC1A5 which is consistent with the report that transformed cells with strong PI3K-Akt-mTOR, KRAS or Myc pathway activation increase their conversion of glutamate to α-ketoglutarate by glutamate dehydrogenase (GLUD) for metabolism and biosynthesis (60–62). Several studies have demonstrated a direct link between hypoxia or hypoxia inducible factor (HIFs) in the development of IPF (63, 64) and It is well recognized that oncogene cMyc enhances glutamine usage by directly transactivating the expression of SLC1A5 in cancer (65, 66). Recently we demonstrated that cMyc is required for TGF-β–induced accumulation of hexokinase 2 in pulmonary fibrosis (67). Here we demonstrated that knockdown of HIF1α, HIF2α and cMyc prevented the increase in SLC1A5 by TGF-β, whereas siRNA-mediated loss of SLC1A5 downregulated the TGF-β induction of HIF1α, HIF2α and cMyc supporting a model of sustained HIF1α, HIF2α and cMyc induction in TGF-β /SLC1A5–dependent differentiation and activation of myofibroblasts. These findings indicated that HIF1α, HIF2α and cMyc plays a distinct role in glutamine metabolism through SLC1A5 to promote metabolic adaptations. Autophagy is a basic cellular homeostatic process important to cell fate decisions under nutrient starved condition. Dysregulation of autophagy impacts numerous human diseases including idiopathic pulmonary fibrosis (68). We observed elevated conversion of LC3-I to LC3-II and beclin1 following V-9302 exposure in primary human lung fibroblast which was not unexpected considering the interrelationship between amino acid withdrawal, regulation of mTOR and autophagy (47). Mitochondria plays a central role in energy metabolism and its dysfunction could induce fibrotic diseases (69). Therefore, understanding the process and mechanism of mitochondrial dysfunction is of great therapeutic value for fibrogenesis. We showed SLC1A5 inhibited fibroblasts have diminished OXOPHOS and glycolysis indicating SLC1A5 is a critical regulator for mitochondrial metabolic reprogramming in fibrosis.

We next ascertained whether the above-mentioned metabolic regulatory network represents a novel approach that can be targeted in a murine treatment model of pulmonary fibrosis. The preclinical efficacy of SLC1A5 inhibitor, V-9302 was assessed in the bleomycin model of lung fibrosis. V-9302 treatment could robustly block glutamine uptake in mice tumor model but had no influence in normal tissue and T cell activation and viability after V-9302 exposure (21, 45). Over the course of V-9302 exposure, liver and kidney histology was similar among C57BL6 male/female mice treated with either V-9302 or vehicle without any demonstrable effect on murine liver enzymes or circulating blood cell populations. As shown in figure 8, not only were profibrotic mediators such as *Col1a1*, *Col3a1, Col4a1, Fn*, *Pai-1* and *Ctgf* inhibited by V-9302, but peripheral blood oxygenation on room air (SpO_2_; an overall indicator of lung physiology that has been shown in various animal models to accurately predict lung function; (70–72) and lung compliance by Flexivent were similarly improved following SLC1A5 inhibition.

**Figure 8.**
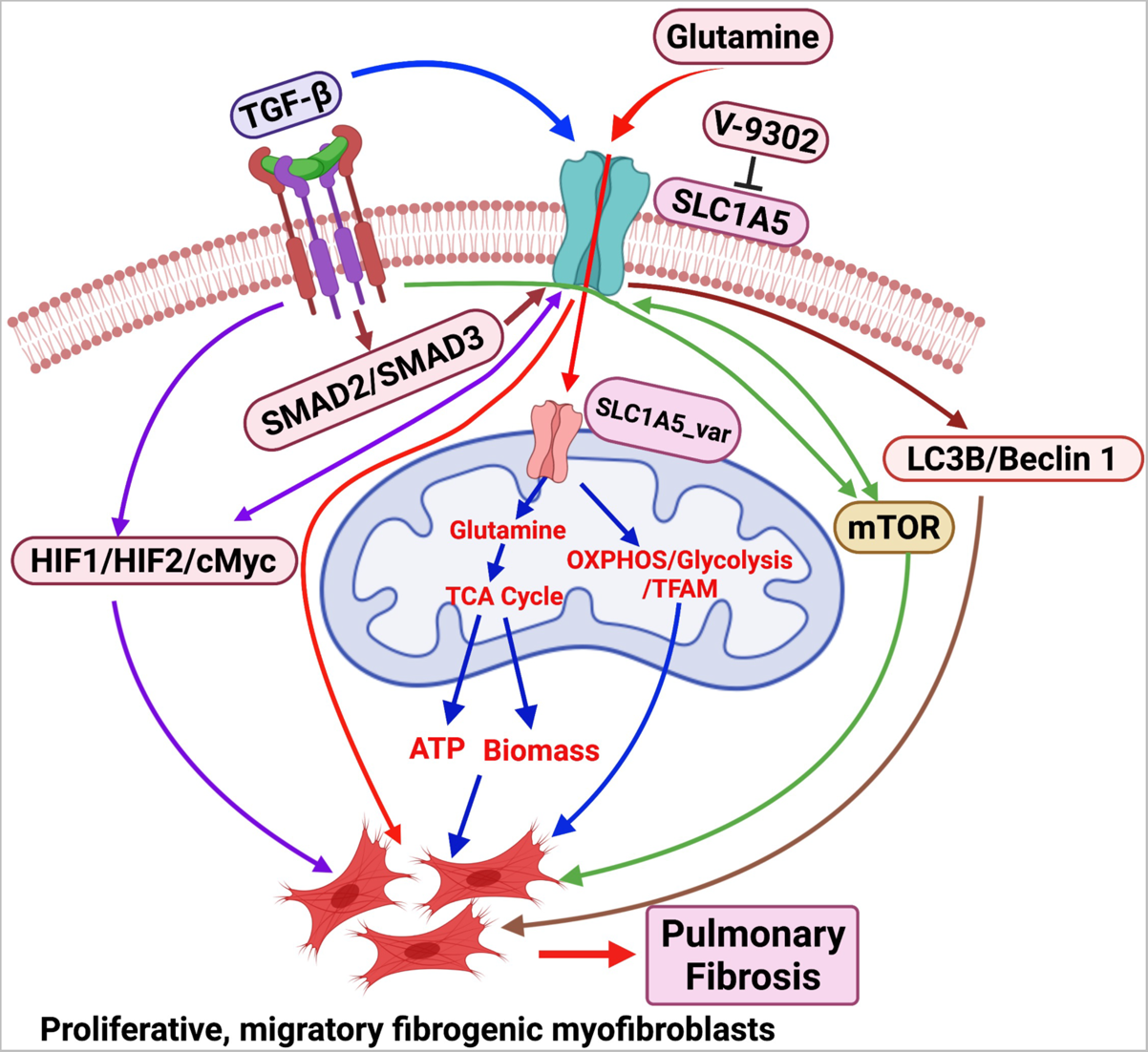
Schematic diagram on the role of SLC1A5 in Pulmonary fibrosis. Glutamine, transported into cells through cell surface glutamine transporter i.e., SLC1A5, to enable ATP production through TCA cycle and to provide nitrogen, sulfur, and carbon as biosynthetic precursors for growing and proliferative fibroblasts. Loss or inhibition of SLC1A5 function enhances the fibroblast susceptibility to autophagy, suppresses mTOR, HIF, Myc signaling and impairs mitochondrial function, ATP production and glycolysis, thus ameliorating pulmonary fibrosis.

A potential limitation in our study is that around 30 amino acid transporters have been identified in human physiology; however, we show that the fibroproliferative actions of TGF-β reflect an extensive metabolic adaptation through a single transporter, SLC1A5. This finding was surprising given that SLC38A1/2/3 were also reported as glutamine transporters upregulated in cancer with transporter plasticity and redundancy (21). Previous work suggested that SLC1A5 deficiency was compensated by SLC38A1 and SLC38A2 in osteosarcoma cells (53). However, in fibroblasts, we did not observe any significant upregulation of other glutamine transporters upon SLC1A5 knock down confirming there was not significant transporter plasticity or redundancy in these cells. While we observed that SLC1A5 significantly alters glucose and fatty acid metabolism (data not shown), we believe further studies will be required to find relevant crosstalk between SLC1A5, glucose, and fatty acid metabolism.

Together our findings demonstrated that glutamine transporter SLC1A5 is a gatekeeper for glutamine metabolism and metabolic reprogramming in pulmonary fibrosis and targeting SLC1A5 could be a new strategy for IPF treatment. This is the first study, to our knowledge, to demonstrate the utility of a pharmacological inhibitor of glutamine transport in fibrosis, representing a potential new class of cell selective targeted therapy and laying a framework for paradigm-shifting treatments targeting cellular metabolism.

## Methods

### Cell culture

Primary human lung fibroblasts (NHLF) were purchased from Lonza and cultured in Dulbecco’s modified Eagle’s medium (DMEM, Life Technologies) supplemented with 10% fetal bovine serum (FBS, Hyclone Laboratories) with the addition of penicillin-Streptomycin (P/S, Life Technologies). De-identified human normal lung fibroblast and IPF fibroblasts, passage no. 3-5, characterized in previously published studies (73, 74), were obtained from Dr. Carol Feghali-Bostwick, Medical University of South Carolina, Charleston, SC, (University of Pittsburgh IRB #970946) and Dr. Nathan Sandbo, University of Wisconsin-Madison, Madison, WI [Translational Science Biocore (TSB) Biobank IRB no. 2011-0521]. They were cultured in 10% FBS/DMEM supplemented with glutamine (Cells from C. Feghali-Bostwick) (73) or no additional additives (Cells from N. Sandbo) (74). Murine embryonic fibroblast cell lines (AKR-2B) (59) were cultured in DMEM supplemented with 10% FBS. BEAS-2B were maintained in RPMI-1640 (Thermo Fisher) + 10% FBS whereas RAW cells were cultured in DMEM supplemented with 10% FBS. All cell lines were maintained at 37^0^C with 5% CO_2_. A summary of pharmacological inhibitors used are listed in table S1.

### Mice

WT male and female C57BL/6 mice were purchased from Charles River Laboratories and used at 10 weeks of age.

### Western blotting

Unless stated otherwise, cells were seeded into six-well plates for 24 hours (2 × 10^5^ cells per well) and changed into medium with 0.1% FBS for 24 hours, and reagents were added for the indicated times. Cells were lysed in ice-cold modified RIPA buffer (50 mM Tris-HCl at pH 8.0, 150 mM NaCl, 1% Triton X-100, 0.1% SDS, 0.5% sodium deoxycholate, 1 mM EDTA (pH 8), 1 mM EGTA, 1 mM PMSF) containing cOmplete Protease Inhibitor mix (Roche), centrifuged at 13,000xg for 15 min at 4^0^C to remove insoluble material and supernatant collected. Equal amounts of each sample (5-20 μg) were vigorously mixed with 5x SDS loading buffer, boiled for 5 min, and separated by 10-12% SDS-polyacrylamide gel electrophoresis. The resolved protein samples were transferred onto Immobilon-P PVDF Membrane (Millipore Sigma), and the blots incubated in the presence of primary antibodies (provided in table S1) at 4^0^C overnight. Immunoreactivity was detected by enhanced chemiluminescence (ECL, Western blotting detection reagents, Thermo Scientific). Quantification was done using Image J software from NIH.

### Measurement of Glutamine

Intracellular glutamine was determined using Glutamine Colorimetric Assay Kit (K556, BioVision) following manufacturer’s instructions. Briefly, NHLF cells were seeded in 6-well plates at a density of 2 x 10^5^ cells/well (10% FBS/DMEM) and incubated for 24h. The medium was changed to 0.1% FBS/DMEM for 24 hours after which cells were pretreated with either dimethyl sulfoxide (DMSO) (0.1%) or SLC1A5 inhibitor V-9302 (10 μM) for 1h and then stimulated with either vehicle [4 mM HCl/0.1% bovine serum albumin (BSA)] or TGF-β (5 ng/ml) for 24 hours and glutamine level determined.

### Real time quantitative PCR (qRT-PCR) analysis

Total RNA was isolated from cells using the RNeasy Plus Mini Kit (Qiagen) and 1 μg RNA reverse-transcribed to cDNA with random primers (Life Technologies) and Maxima Reverse Transcriptase (Thermo Fisher Scientific). The cDNAs (1-5 ng RNA equivalents) were used for real-time PCR amplification using SYBR Premix Ex Taq II (Takara/Clontech) on a 7500 Fast Real-Time PCR System (Applied Biosystems). TATA box binding protein (TBP) or SMAD4 was used as a normalization control. The relative expression levels of the target genes were determined by the 2^-ΔΔ^*^Ct^* method (75). A summary of human and murine primer sets used were listed in table S2. All experiments were performed in triplicate.

To extract RNA from mouse lung, tissues were lysed and homogenized with RLT Plus Buffer (RNeasy Plus Mini Kit; Qiagen). After the lysate passed through a gDNA Eliminator spin column, ethanol was added, and the samples were applied to a RNeasy Min Elute spin column according to the manufacturer’s instructions.

### siRNA mediated gene knockdown

NHLF, IPF fibroblast or murine AKR-2B cells were transiently transfected with 40 nM of small interfering RNA (siRNA) to SLC1A5 (sc-60210 or sc-60211), HIF-1α (sc-35561), HIF-2α (sc-35316), cMyc (sc-29226), SMAD2 (sc-38374), SMAD3 (sc-38376), mtTFA (sc-38053) or non-targeting control (NT; sc-37007) (Santa Cruz Biotechnology) according to the manufacturer’s protocol. siRNA from Santa Cruz contains a pool of 3 target-specific siRNAs for murine or human cells. In brief, 7.5×10^4^ cells were transfected using Lipofectamine 3000 (Invitrogen) and incubated in Opti-Mem (Invitrogen) for 6h. Cells were then placed in complete medium (10% FBS/DMEM) for 18h to recover prior to addition of Vehicle or TGF-β in low serum medium (0.1% FBS/DMEM) for the indicated times and processed for Western blot or qPCR analysis.

### Rescue SLC1A5 inhibition

NHLF cells were transfected with siCt or siSLC1A5 using Lipofectamine 3000 for 24h. After treating with either water (vehicle) or 2 mM diethyl-2-oxopentanedioate (esterified form of a-KG) for 24h in 0.1% FBS/DMEM, cultures were stimulated with vehicle or TGF-β (5 ng/ml) for another 24 hours at 37°C. Cultures were then processed for Western blotting.

### Scratch assay

For scratch assays, 2×10^5^ NHLF or 1.5×10^5^ AKR-2B cells were seeded into 6-well plates containing 10% FBS/DMEM. Following 24h of incubation, the monolayer was scraped in a straight line to create a “scratch” with a sterile p200 pipet tip. Cultures were then treated with V-9302 or GPNA or the indicated siRNA and incubated in the presence or absence of TGF-β (5 ng/ml) for 24 hours at 37°C. Images were taken at 24h and the cellular leading edge was quantitated using Image J software from NIH (76).

### Soft agar colony formation assay

Soft agar assays were performed as previously described (77). Briefly, 1.25×10^4^ cells were seeded into 6-well plates in the presence or absence of TGF-β (25 ng/ml) ± V-9302 or GPNA. After 7 d growth (peak colony formation) at 37°C, colonies ≥ 50 μm in diameter were counted using an optimized CHARM algorithm in Optronix Gelcount (Oxford Optronics). All experiments were performed in triplicate.

### MTT assay

The cellular toxicity of V-9302 was measured by MTT assay. Briefly, 2.5×10^3^ NHLF cells were seeded into 96 well plates with 10% FBS/DMEM and incubated overnight. Cells were then treated with the indicated concentration of V-9302 (1-40 μM) in 10% FBS/DMEM or 0.1% FBS/DMEM in a total volume of 100 μl. Following 24 h incubation, MTT (10 μl of 5 mg/ml MTT in PBS) was added and incubated for another 4 h at 37°C. After the incubation, medium was removed and 100 μl DMSO was added for 10 min and the color reaction measured at 570 nm using a microplate reader.

### XTT Assay for Cell Viability and Proliferation

XTT is a colorimetric assay for the nonradioactive quantification of cellular proliferation, viability, and cytotoxicity. Briefly, NHLF cells were seeded into a 96 well microplate at a concentration of 2.5×10^3^ cells/well in 100 μl of 10% FBS/DMEM and incubated overnight. Cells were then treated with various amounts of V-9302 (1-40 μM) in 10% FBS/DMEM and Incubated for 24h at 37°C and 5% CO_2_. After 24h incubation, 50 µl of XTT labeling mixture were added per well and incubated for another 4h. Spectrophotometrical absorbance of the formazan product were measured using a microplate (ELISA) reader at 450 nm.

### Seahorse assay

NHLF (2 × 10^5^ per well) cells were seeded into six-well plates for 24h (10% FBS/DMEM). Following 24h of serum starvation (0.1% FBS/DMEM), cells were pretreated with DMSO (0.1%) or V-9302 (10 µM) at 37°C for 1 hour followed by vehicle or TGF-β (5 ng/ml) and 3000 cells were seeded into Seahorse assay microplates (Agilent Technologies). After 24h of incubation, the medium was changed to Seahorse XF base assay medium supplemented with 1 mM sodium pyruvate, 2 mM l-glutamine, and 10 mM glucose (OCR) or with 2 mM glutamine (ECAR), adjusted to pH 7.4 and incubated for 1h at 37°C in a non-CO_2_ incubator to equilibrate the CO_2_ level in the atmosphere. OCR and ECAR was measured on a Seahorse XFp extracellular flux analyzer (Seahorse Bioscience, Agilent Technologies) as described previously (67) by using the Seahorse XFp Cell Mito Stress Test Kit and XF glycolysis stress test kit (Agilent Technologies). OCR and ECAR were measured under basal conditions and in response to several metabolic drugs provided in the kit, such as oligomycin (1 µM), FCCP (carbonyl cyanide-p-trifluorome-thoxyphenylhydrazone, 0.5 µM), Rotenone/Antimycin A (0.5 µM /0.5 µM), glucose, (25mM), and 2-DG (2-deoxyglucose, 50 mM).

### Immunofluorescence (Acta2) staining

AKR-2B cells (1×10^5^) in 10% FBS/DMEM were incubated onto coverslips in 6-well plates for 24h. After 24h incubation, medium was replaced with 0.1% FBS/DMEM, treated with either Vehicle (4 mM HCl/0.1% BSA) or TGF-β (5 ng/ml) with or without GPNA (10 mM) for 24h. After 24h incubation, cells were fixed with 4% paraformaldehyde, permeabilized with 0.1% TritonX-100, blocked with blocking buffer (5% Normal goat serum, 1% Glycerol, 0.1% BSA, 0.1% Fish skin Gelatin, 0.04% Sodium Azide, PBS pH 7.2) at RT for 1h and incubated with Acta2 antibody (Sigma) at 4^0^C overnight. After 3X PBS washing, Acta2 was labeled with Alexa Fluor® 594 goat anti-mouse secondary antibody (Life technologies) at RT for 30m with DAPI. Fluorescence images were collected on an LSM510 confocal microscope (Carl Zeiss Microimage Inc.), and density analyzed by ImageJ software from NIH.

### Bleomycin model of pulmonary fibrosis

Ten-week-old male and female C57BL/6 mice were administered bleomycin (BLM; 3.5 U/Kg) or 75 μl of 0.9% normal saline alone by tracheal instillation using an intratracheal aerosolizer (Penn-Century, Inc., Wyndmoor, PA) on day 0, while under ketamine/xylazine anesthesia. On day 11, BLM-treated mice were randomly assigned to receive V-9302 as a soluble intraperitoneal (IP) injection (0.2 ml) in PBS supplemented with 2% DMSO daily until day 24. AS V-9302 concentration of 50-75 mg/kg/day has been shown to be optimal in various tumor models (21, 28), we used 37.5 mg/kg/day of V-9302. Animals were shaved around the collar region to allow determination of dissolved oxygen (SpO2) levels using the MouseOx monitoring system (Starr Life Science) on day 3, 6, 9, 13, 16, 21. On day 25, mice were given pentobarbital (100 mg/kg) and a small incision was made in the neck to expose the trachea. A metal cannula was placed between the rings of the trachea and secured in place with a suture. The mouse was connected to the ventilator and measurements for static and dynamic compliance was recorded. Following death, lung was excised, weighed, and collected for hydroxyproline, histopathology, and fibrotic marker expression determination.

### Hydroxyproline assay

Total lung collagen levels were assessed using the Hydroxyproline Assay Kit (MAK008, Sigma-Aldrich). Briefly, 20 mg of mouse lungs were homogenized in H_2_O to 100 mg/ml concentrations. 100 μl of homogenates were mixed with 100 μl HCl (12M) and hydrolyzed at 120^0^C overnight. 10 μl of supernatants were then analyzed to a 96-well plate for hydroxyproline according to the manufacturer’s instructions.

### Histological Scoring

All H&E-stained slides were reviewed by a pulmonary pathologist. Blinded histologic scoring assay was used in determining development and resolution of fibrosis. Semiquantitative evaluation considering both intensity and extent were performed, and scores added for fibrosis. Scoring was performed based on the following criteria: 0 (no fibrosis), 1 (focal/minimal fibrosis), 2 (multifocal/moderate fibrosis), 3 (confluent/severe fibrosis).

### Statistics

Statistical significances were calculated by using GraphPad Prism 9.3 software. All in vitro data were from a minimum of 3 biological replicates. In vivo data represented 3 male and 3 female mice per group. Results shown reflect mean ± SEM. The difference between 2 groups was analyzed using a two-tailed unpaired student’s t test. When more than 2 groups were used, statistical analysis was performed by either one-way or two-way ANOVA followed by Tukey’s post hoc test. Statistical significance is denoted by the number of asterisks, *P < 0.05, **P < 0.01, ***P < 0.001, ****P < 0.0001. A p value lower than 0.05 was considered significant.

### Study Approval

All studies involving mice were performed according to Mayo Clinic approved IACUC protocol #A00005741-20. All human lung fibroblasts studies were approved previously by the University of Pittsburgh (IRB #970946) or University of Wisconsin-Madison (IRB #2011-0521) Ethics Committee.

## Author contributions

M.C., D.J.T., and A.H.L., designed research; M.C., K.J.S., and T.J.K. performed research, with the majority done by M.C; E.S.Y., analyzed histology of mouse lung; M.C., D.J.T., and A.H.L., wrote the manuscript. All authors analyzed data and edited the manuscript.

## Acknowledgements

We would like to thank Dr. Carol Feghali-Bostwick (Medical University of South Carolina, Charleston, SC) and Dr. Nathan Sandbo (University of Wisconsin-Madison, Madison, WI) for providing primary lung fibroblasts from healthy and IPF patients. This work was supported by grants from the National Institutes of Health (NIH Contract 268201100020C-7-0-1), Robert N. Brewer Family Foundation and Caerus Foundation (A.H.L.). The research received support from the “Brewer Family Career Development Award in Support of Idiopathic Pulmonary Fibrosis and Related Interstitial Lung Disease Research” and “Boehringer Ingelheim Discovery Award in Idiopathic Pulmonary Fibrosis/Interstitial Lung Disease” (M.G.).

