## Supplemental File for "Targeting Pulmonary Fibrosis by SLC1A5 dependent Glutamine Transport Blockade"

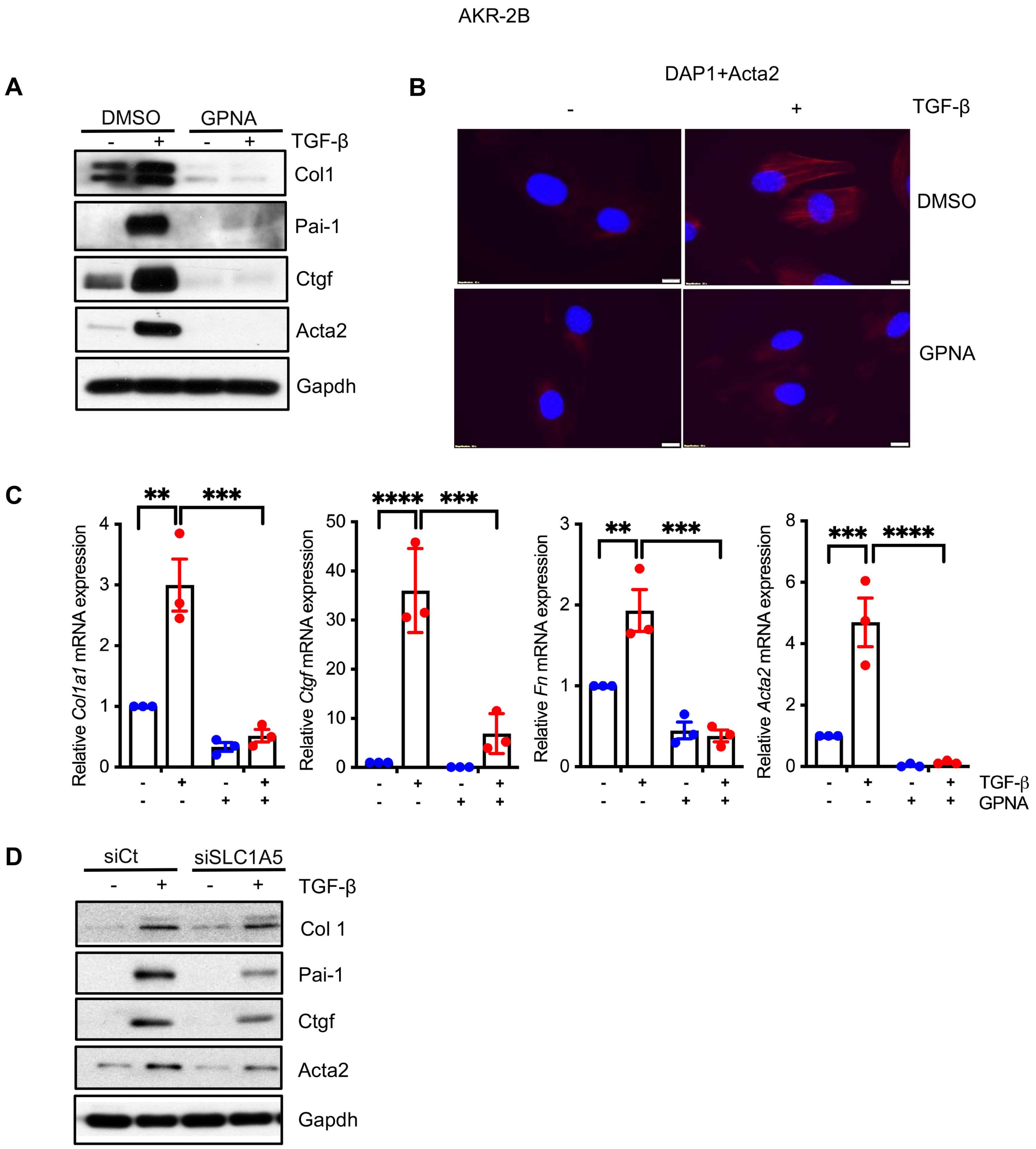


**Supplementary Figure 1. Profibrotic TGF-β signaling is dependent on the action of SLC1A5**. Mouse embryonic fibroblast (AKR-2B) cells were treated with GPNA (10 mM) for 1h (**A, B, C**) or transfected with non-targeting control (siCt) or siRNA against SLC1A5 (siSLC1A5) (**D**). Vehicle (-) or TGF-β (+) was directly added to a final concentration of 5 ng/ml and following 48 h incubation, lysates were prepared and Western blotted for Col1, Pai-1, Ctgf, Acta2 and Fn. GAPDH was used as a loading control (A, D), (**B**) Immunofluorescence staining of Acta2 in AKR-2B cells with/without GPNA (10 mM). (**C**) qPCR analysis of indicated profibrotic genes. n = 3. Differences between groups were determined by two-way ANOVA test with Tukey post-hoc analysis (C) using GraphPad Prism 9.3 software. **P < 0.01, ***P < 0.001, ****P < 0.0001.

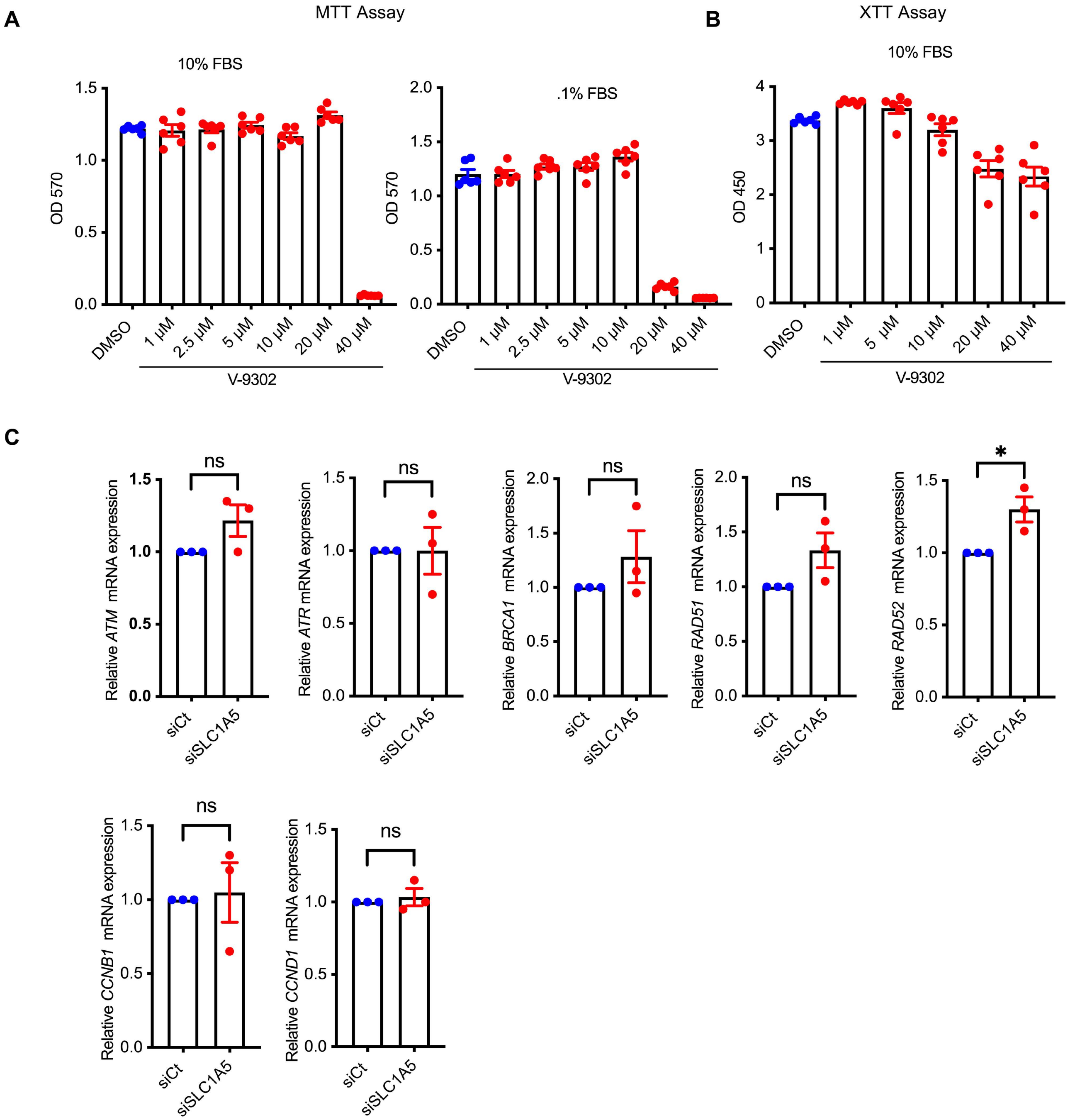


**Supplementary Figure 2. SLC1A5 inhibitor V-9302 does not inhibit in vitro cell proliferation and viability, DNA repair, recombination and cell cycle regulated gene expression**. (**A, B**) NHLF cells (DMEM/10% FBS) were seeded at 2.5x$10^3$ cells/96 well plate. 24h after seeding, the medium was removed and replaced with either 10%FBS/DMEM or 0.1% FBS/DMEM containing vehicle (0.1% DMSO) or V-9302 (0,1, 2.5, 5, 10, 20 and 40 µM) for 24 h prior to MTT and XTT assay. Data reflect mean ± SEM for n = 6. (**C**) Expression of DNA repair (*ATM, ATR and BRCA1*), recombination (*RAD51* and *RAD52*) and cell cycle regulated genes (*CCNB1* and *CCND1*) determined by qPCR in proliferating SLC1A5 knocked down NHLF cells. Data represent mean ± SEM for n = 3.

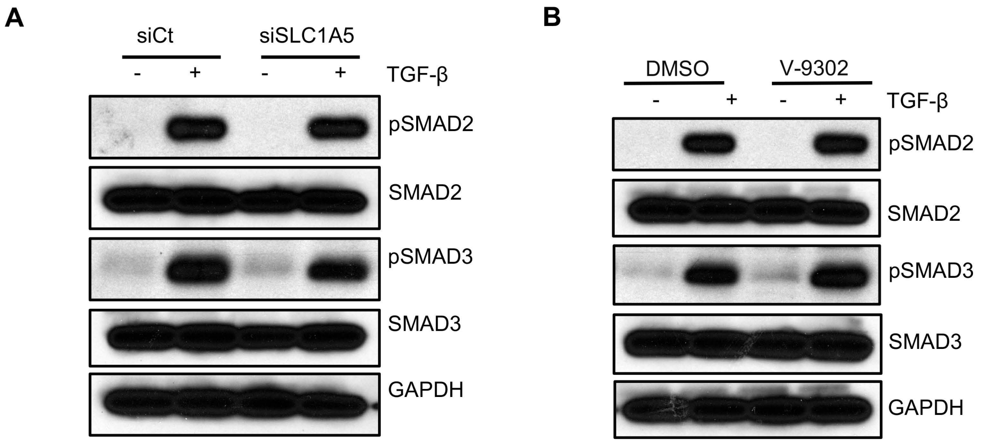


**Supplementary Figure 3**. **Phosphorylation of SMAD2 or SMAD3 occurs independently of SLC1A5**. NHLF cells were transfected with scrambled control or SLC1A5 siRNA (**A**) or treated with V-9302 (**B**) and stimulated in the absence (-) or presence (+) of TGF-β (5 ng/ml) for 6h. Western blotting of harvested cell lysates was performed for pSMAD2, SMAD2, pSMAD3 and SMAD3. GAPDH was used as loading control. Data are representative of 3 separate experiments.

A

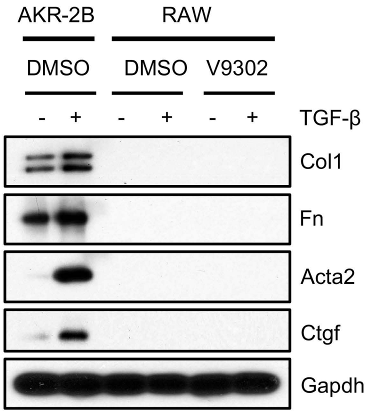


B

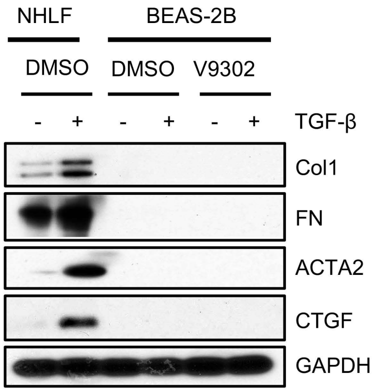


**Supplementary Figure 4. Profibrotic protein expression in epithelial and alveolar macrophages are not influenced by TGF-β or V-9302**. AKR-2B (murine fibroblast), and RAW (murine macrophages) (**A**), or NHLF (human lung fibroblast) and BEAS-2B (human epithelial) (**B**) cells were treated with V-9302 (10 μM), stimulated in the absence (-) or presence (+) of TGF-β (5 ng/ml) for 48h and Western blotted for the profibrotic molecules.

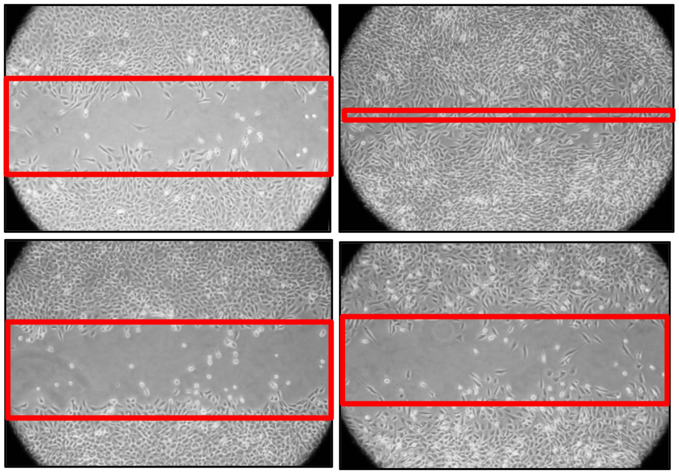


**Supplementary Figure 5. Cell migration is dependent on SLC1A5 activity.** Scratch assays were performed on AKR-2B cells. Red bands indicate the leading edge following 24h in the presence and absence of TGF-β with or without GPNA (10mM).

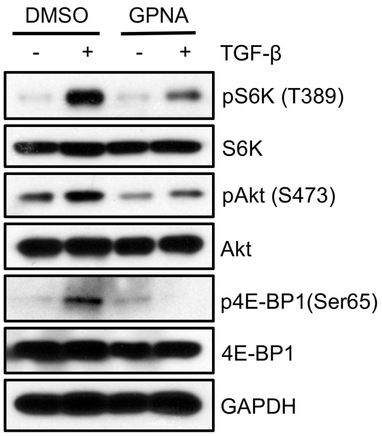


**Supplementary Figure 6. Role of SLC1A5 in mTOR signaling**. AKR-2B cells were pretreated with GPNA (10mM for 1h) and Western blotted for the indicative proteins 6h post TGF-β or vehicle treatment.

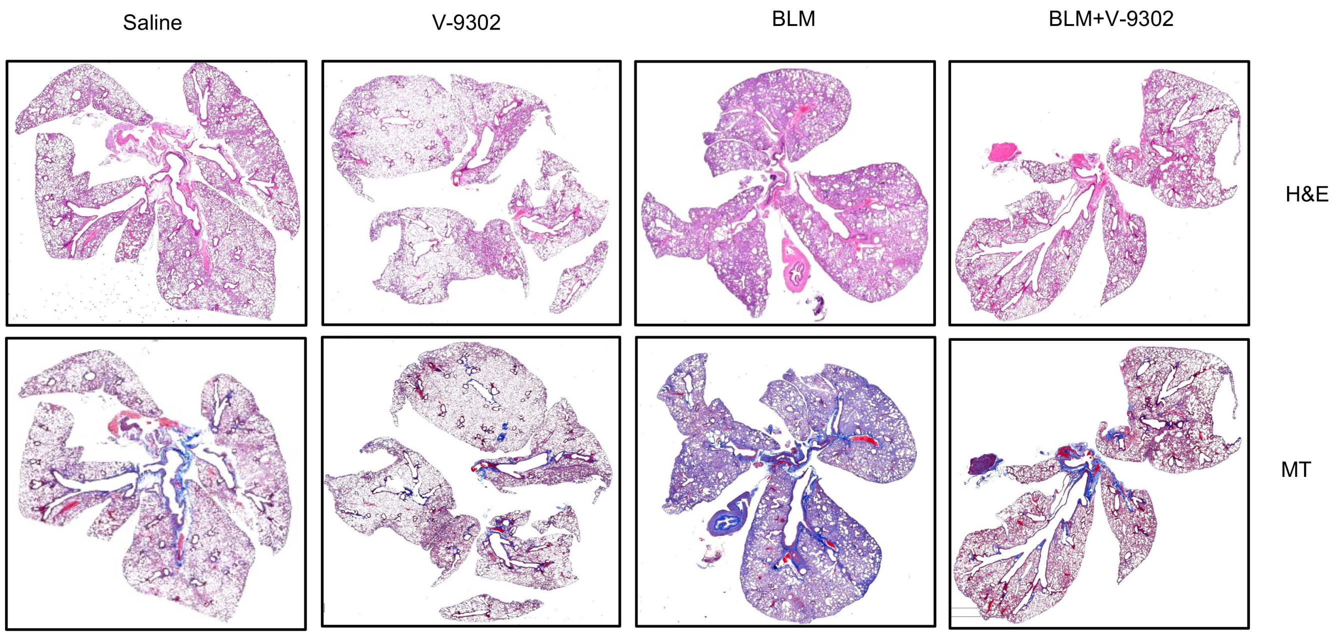


**Supplementary Figure 7**. **V-9302 attenuates bleomycin induced pulmonary fibrosis on lung structure**. (A) Mice were treated as in Figure 7A, and Hematoxylin and Eosin (H&E) staining for histology and Masson's trichrome (MT) for collagen deposition (blue) were performed following euthanasia at day 25. Representative images from 6 mice are shown. Scale bars, 3 mm.

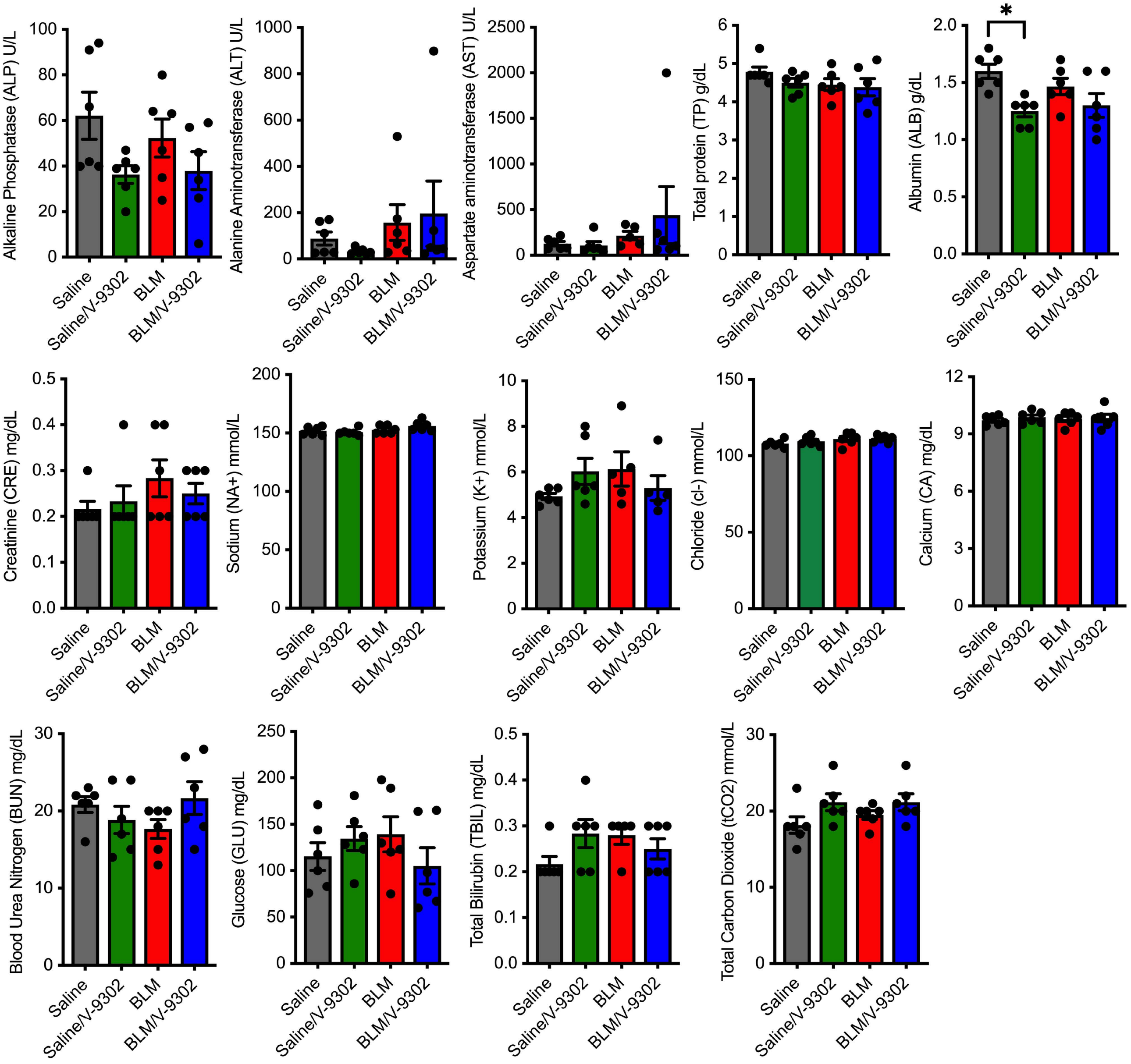


**Supplementary Figure 8**. **V-9302 has no demonstrable effect on murine liver or kidney**. C57BL/6 male/female mice were treated as described in Methods and Figure 7A. (**A**) On day 25, blood samples were collected in lithium heparin tube from the facial vein of unanesthetized animals. Serum levels of indicated parameters were determined using a Piccolo Xpress Chemistry Analyzer. The Toxicology and Pharmacology Laboratory in the Department of Molecular Medicine at the Mayo Clinic performed all analyses. Data are presented as mean ± SEM of n = 6.

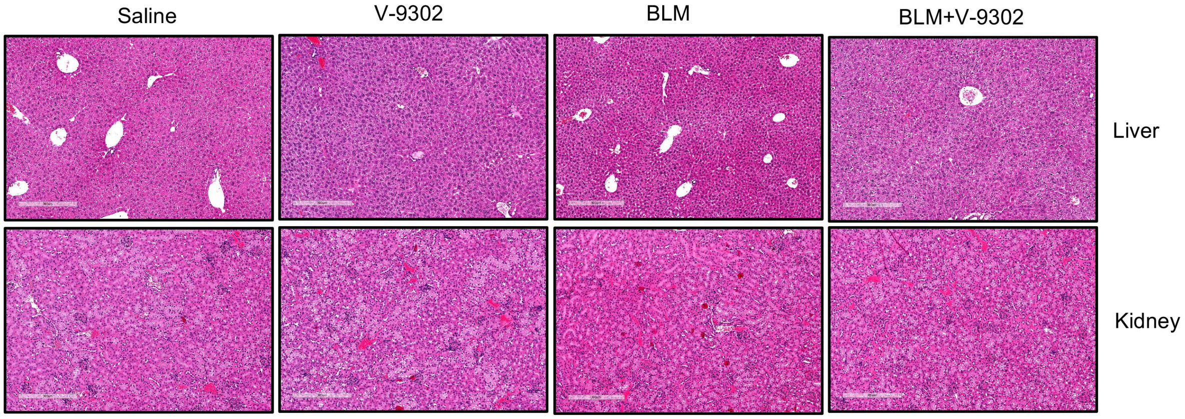


**Supplementary Figure 9. Liver and kidney pathology was similar among mice chronically treated with either V-9302 or vehicle**. Hematoxylin and Eosin (H&E) staining of liver and kidney histology. Data are presented as of n = 6 mice. Scale bars 300 μm.

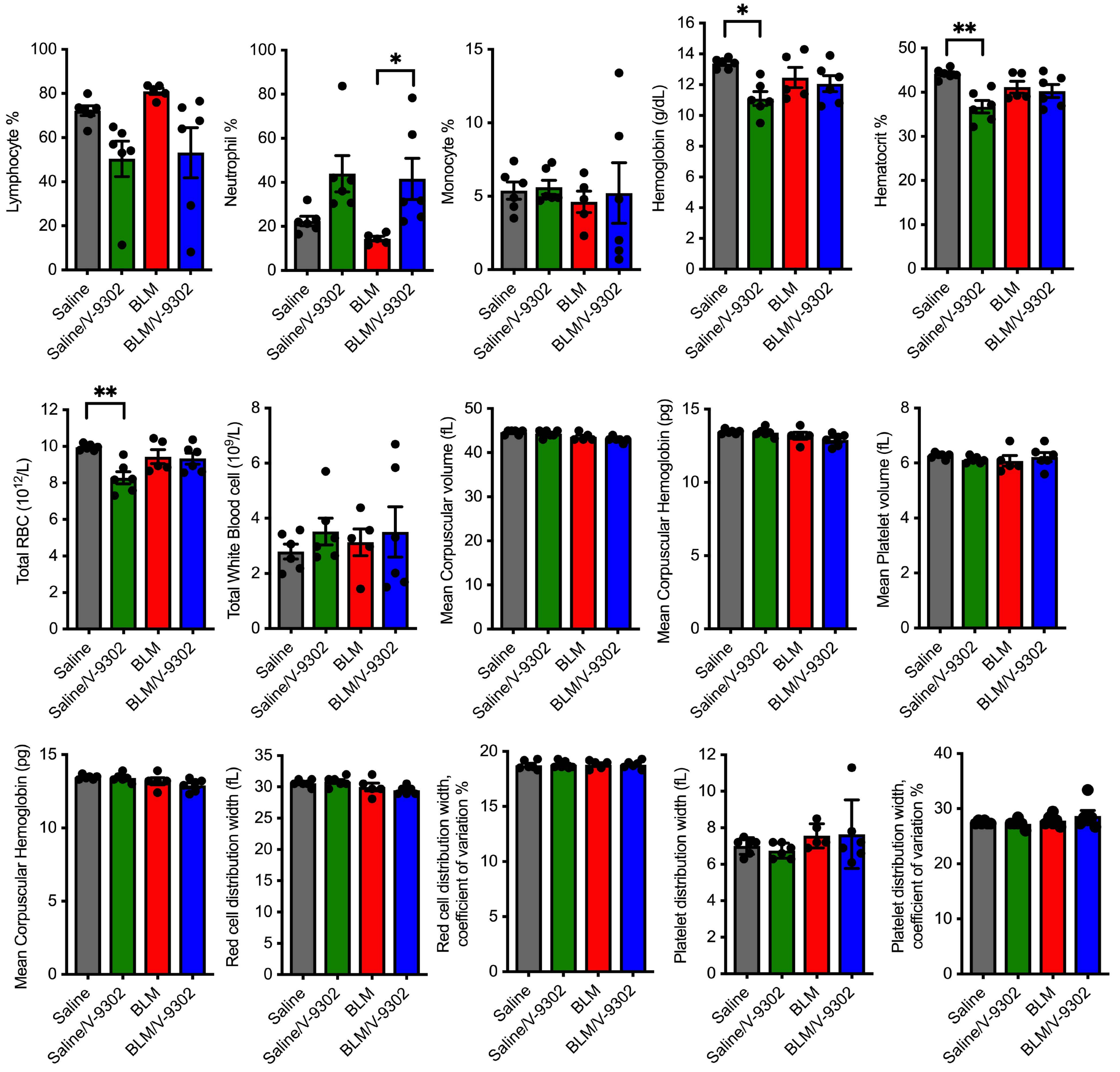


**Supplementary Figure 10. V-9302 treatment does not impair inflammatory cell recruitment.** C57BL/6 male/female mice were treated as described in Methods and Figure 7A. On day 25, blood samples were collected in EDTA. Quantification of inflammatory cells and other blood parameters were conducted using a VetScan HM5 Analyzer at The Toxicology and Pharmacology Laboratory in the Department of Molecular Medicine of Mayo Clinic. Data are represented as mean ± SEM of n = 6.

**Table S1**

| REAGENT or RESOURCE | SOURCE | IDENTIFIER |
|---|---|---|
| Antibodies | | |
| Rabbit polyclonal anti-SLC1A5 Antibody | Cell Signaling Technology | Cat# #5345S |
| Rabbit polyclonal anti-SLC1A5 Antibody | Cell Signaling Technology | Cat# 8057S |
| Mouse monoclonal Anti- GAPDH antibody | Millipore | Cat# MAB374 |
| Goat polyclonal anti-Type I Collagen antibody | Southern Biotech | Cat# 1310-01 |
| Goat polyclonal anti-PAI-1 antibody | R & D Systems | Cat# AF3828 |
| Goat polyclonal anti-CTGF antibody | Santa Cruz Biotechnology | Cat# sc-14939 |
| Rabbit monoclonal anti-CTGF antibody | Cell Signaling Technology | Cat# 86641S |
| Rabbit polyclonal anti-phospho-SMAD3 antibody | (1) | N/A |
| Rabbit monoclonal anti-SMAD3 antibody | Abcam | Cat# ab40854 |
| Mouse monoclonal anti-αSMA antibody | Sigma-Aldrich | Cat# A5228 |
| Rabbit polyclonal anti-Fibronectin antibody | Sigma-Aldrich | Cat# F3648 |
| Rabbit polyclonal anti-SMAD2 antibody | Abcam | Cat# ab63576 |
| Rabbit polyclonal anti-phospho-Smad2 antibody | (1) | N/A |
| Rabbit polyclonal anti-Akt antibody | Cell Signaling Technology | Cat# 9272 |
| Rabbit polyclonal anti-phospho-Akt (Ser473) antibody | Cell Signaling Technology | Cat# 9271 |
| Rabbit polyclonal anti-p70 S6 Kinase antibody | Cell Signaling Technology | Cat# 9202 |
| Rabbit polyclonal anti-phospho-p70 S6 Kinase (Thr389) antibody | Cell Signaling Technology | Cat# 9205 |
| Rabbit monoclonal anti-Phospho-4E-BP1 antibody | Cell Signaling Technology | Cat# 2855S |
| Rabbit monoclonal anti-4E-BP1 antibody | Cell Signaling Technology | Cat# 9644S |
| Rabbit polyclonal anti-LC3B antibody | Abcam | Cat#ab48394 |
| Rabbit polyclonal anti-Beclin 1 antibody | Abcam | Cat#ab62557 |
| Rabbit monoclonal anti-ATG7 antibody | Abcam | Cat#ab133528 |
| Rabbit monoclonal anti-ATG5 antibody | Abcam | Cat#ab108327 |
| Rabbit monoclonal anti-HIF-1α antibody | Cell Signaling Technology | Cat# 14179S |
| Rabbit monoclonal anti-HIF-2α antibody | Cell Signaling Technology | Cat# 7096S |
| Rabbit monoclonal anti-c-Myc antibody | Cell Signaling Technology | Cat# 5605S |
| Biological Samples | | |
| Normal and IPF lung fibroblast | This manuscript and (2) | U of Pittsburgh IRB#970946 |
| Chemicals, Peptides, and Recombinant Proteins | | |
| DMEM (Dulbecco's Modified Eagle Medium) | Life Technologies | Cat#11965-092 |
| RPMI 1640 | Thermo Fisher | Cat# 11875093 |

| | | |
|---|---|---|
| Fetal Bovine Serum | HyClone Laboratories | Cat#SH3007103 |
| Penicillin-Streptomycin | Life Technologies | Cat#15070063 |
| L-Glutamine | Life Technologies | Cat#25030081 |
| Protease Inhibitor Cocktail | Roche | Cat#11836153001 |
| V-9302 | Selleckchem | Cat# S8818 |
| L-Glutamic acid γ-(p-nitroanilide) hydrochloride (GPNA) | Sigma | Cat# G6133 |
| LY294002 | Sigma-Aldrich | Cat#L9908 |
| MK2206 | Selleckchem | Cat#S1078 |
| Rapamycin | Selleckchem | Cat#S1039 |
| SB431542 | Tocris | Cat#161410 |
| Lipofectamine 3000 | Thermo Fisher | Cat# L3000001 |
| U0126 | Promega | Cat#V1121 |
| Torin 1 | Selleckchem | Cat#S2827 |
| Rotenone | Selleckchem | Cat#S2348 |
| 3-Nitropropionic acid | Selleckchem | Cat# S3652 |
| Antimycin A | Sigma | Cat# A8674 |
| Thiazolyl Blue Tetrazolium Bromide | Sigma | Cat# M5655 |
| Critical Commercial Assays | | |
| Glutamine Colorimetric Assay Kit | BioVision | Cat# K556 |
| RNeasy Plus Mini Kit | Qiagen | Cat#74136 |
| TB Green® Premix Ex Taq™ II (Tli RNase H Plus) | Takara | Cat#RR820B |
| Maxima Reverse Transcriptase | Life Technologies | Cat#EP0741 |
| Hydroxyproline Assay Kit | Sigma-Aldrich | Cat#MAK008 |
| Agilent Seahorse XF Cell Mito Stress Test Kit | Agilent Technologies | Cat#103015-100 |
| Agilent Seahorse XF Glycolysis Stress Test Kit | Agilent Technologies | Cat#103020-100 |
| ATP Colorimetric/Fluorometric Assay Kit | BioVision | Cat# K354 |
| Cell Proliferation Kit II (XTT) | Roche | Cat# 11 465 015 001 |
| Experimental Models: Cell Lines | | |
| Primary Human Lung fibroblast (NHLF) | Lonza | Cat#CC-2512 |
| AKR-2B | (3) | N/A |
| BEAS-2B | ATCC | Cat#CRL-9609 |
| RAW | ATCC | Cat#TIB-71 |
| Oligonucleotides | | |
| siRNA targeting human SLC1A5 | Santa Cruz Biotechnology | Cat#sc-60210 |
| siRNA targeting human HIF1α | Santa Cruz Biotechnology | Cat#sc-35561 |
| siRNA targeting human HIF2α | Santa Cruz Biotechnology | Cat# sc-35316 |
| siRNA targeting human c-Myc | Santa Cruz Biotechnology | Cat# sc-29226 |
| siRNA targeting human SMAD2 | Santa Cruz Biotechnology | Cat# sc-38374 |
| siRNA targeting human SMAD3 | Santa Cruz Biotechnology | Cat# sc-3837 |
| siRNA targeting human mtTFA | Santa Cruz Biotechnology | Cat# sc-38053 |
| siRNA targeting murine SLC1A5 | Santa Cruz Biotechnology | Cat# sc-60211 |

| | | |
|---|---|---|
| Control siRNA | Santa Cruz Biotechnology | Cat#sc-37007 |
| Primers for all quantitative reverse transcription polymerase chain reaction (RT-qPCR), please refer to the Table S2 | This paper | N/A |
| Software and Algorithms | | |
| ImageJ | (4) | https://imagej.nih.gov/ij/ |
| GraphPad Prism 9.3.1 | | https://www.graphpad.com/scientific-software/prism/ |
| BioRender | | https://biorender.com |

Table S2. Primer for human qRT-PCR

| Gene | Forward | Reverse |
|---|---|---|
| SLC1A5 | CTCGATTCGTTCCTGGATCTT | GTTCCGGTGATATTCCTCTCTTC |
| SLC38A1 | AATGCAATGGTGGGTAAAGC | TGACACGTGTACGCCAAAAT |
| SLC38A2 | GCTGCAGATGCACCAATAAA | TTACTTGTCATCTTTGTCCCAAC |
| SLC38A3 | ATCGGAGCCATGTCCAGCTA | AGGGGCAGAATGATGGTGAC |
| SLC38A4 | GAAATTCCAAATACCCTGCCCT | GCGGTGGGTGTAATCCATCA |
| SLC38A5 | GAGTTGCGGCCACTTCAG | TCCATTCATCTTTGGATCCTG |
| SLC7A5 | CCGTGCCGTCCCTCGTGTTC | GGTTCACCTTGATGGGCCGCT |
| SLC7A8 | CTCCACTGGAAAAAAGGTAGCA | TGGTGAATGAAGCCACATCTG |
| SLC7A11 | TGGACGGTGTGTGGGGTCCT | CAGCAGTAGCTGCAGGGCGTA |
| LC3BII | GATGTCCGACTTATTCGAGAGC | TTGAGCTGTAAGCGCCTTCTA |
| ATF4 | TTAAGCCATGGCGCTTCTCA | GCTGGAATCGAGGAATGTGC |
| TRIB3 | TGCCCTACAGGCACTGAGTA | GTCCGAGTGAAAAAGGCGTA |
| PSPH | CACGGTCATCAGAGAAGAAG | GGTTGCTCTGCTATGAGTCT |
| ASNS | GACAGAAGGATTGGCTGCCT | CATCCAGAGCCTGAATGCCT |
| GCN2 | TGCCAACTTACATCAGAAAAGC | TTTGAGGTATATTTGCTTTGG |
| GOT2 | GAGAACAGCGAAGTCTTGAAGAGTG | CTGGCTCCGATCCTTAAGGCT |
| CHAC1 | GGTGACGCTCCTTGAAGATCA | TCAGTGGTTGGTCAGGAGCAT |
| COL1A1 | GAGGGCCAAGACGAAGACATC | CAGATCACGTCATCGCACAAC |
| CTGF | GTCCAGCACGAGGCTCA | TCGCCTTCGTGGTCCTC |
| FN | TGTCAGTCAAAGCAAGCCCG | TTAGGACGCTCATAAGTGTCACCC |
| ACTA2 | GACAATGGCTCTGGGCTCTGTAA | CTGTGCTTCGTCACCCACGTA |
| TGM2 | TCAGCTACAATGGGATCTTGG | AAGGCAGTCACGGTATTTCTC |
| LOX | ACATTCGCTACACAGGACATC | TTCCCACTTCAGAACACCAG |
| LOXL1 | TGCCAGTGGATCGACATAAC | GAAACGTAGCGACCTGTGTAG |
| LOXL2 | GTGCAGCGACAAAAGGATTC | GCGGTAGGTTGAGAGGATG |
| LOXL3 | AGCGAAAAGAGGGTCAACG | TGTCATTGGCACGATAGAACTC |
| LOXL4 | GTGGCAGAGTCAGATTTCTCC | TTGTTCCTGAGACGCTGTTC |
| PLAU1 | GGGAGATGAAGTTTGAGGTGG | AGATGGTCTGTATAGTCCGGG |
| CTSK | CTCCTTCCAGTTTTACAGCAAAG | TTTCCCCAGTTTTCTCCCC |
| MMP14 | TGCCTACCGACAAGATTGATG | ATCCCTTCCCAGACTTTGATG |
| SLC1A5_var | GCCCTCCCACTATGTACTCTA | CTACCAAGCCCAGGATGTTC |
| HIF1$\alpha$ | GTCTGCAACATGGAAGGTATTG | GCAGGTCATAGGTGGTTTCT |
| HIF2$\alpha$ | GACTTACACAGGTGGAGCTAAC | GAGACTCAGGTTCTCACGAATC |

| | | |
|---|---|---|
| c-Myc | CTCCACACATCAGCACAACTA | TGTCCAACTTGACCCTCTTG |
| ATM | CAGGCGAAAAGAATCTGGGG | GCACAAAGTAGGGTGGGAAAGC |
| ATR | TGAAAGGGCATTCCAAAGCG | CAATAGATAACGGCAGTCCTGTCAC |
| BRCA1 | GCAGAGAGTCAGACCCTTCAATGG | GCCCAGGTTTCAAGTTTCCTTTTC |
| RAD51 | CAACCCATTTCACGGTTAGAGC | TTCTTTGGCGCATAGGCAACA |
| RAD52 | GTAGGGAGAGGCTCTGGACA | GCAGGTGCTTAGGACCAAGT |
| CCNB1 | GACCTGTGTCAGGCTTTCTCTG | GGTATTTTGGTCTGACTGCTTGC |
| CCND1 | AGTTGTTGGGGCTCCTCAG | AGACCTTCGTTGCCCTCTGT |
| TBP | GCCCGAAACGCCGAATATAATC | GTCTGGACTGTTCTTCACTCTTGG |
| TFAM | AGCTCAGAACCCAGATGC | CCACTCCGCCCTATAAGC |

Primer for mouse qRT-PCR

| Gene | Forward | Reverse |
|---|---|---|
| Slc1a5 | TGGCCAGCAAGATTGTGGAGAT | TTTGCGGGTGAAGAGGAAGT |
| Col1a1 | ATCTCCTGGTGCTGATGGAC | ACCTTGTTTGCCAGGTTCAC |
| Col3a1 | AGGCAACAGTGGTTCTCCTG | GACCTCGTGCTCCAGTTAGC |
| Col4a1 | CACCCATCTCTGGGGACAAC | TTAGGGCACTGCGGAATCTG |
| Col4a2 | CGGCGTAATCTCAAAAGGCG | GGCCTCTGCTTCCTTTCTGT |
| Pai-1 | TTCCAACCAGCATCCCAGAC | CCATGAGACCTTTGTGGGGT |
| Ctgf | CACAGAGTGGAGCGCCTGTTC | GATGCACTTTTTGCCCTTCTTAATG |
| Fn | TGACAACTGCCGTAGACCTG | ATCTAGCGGCATGAAGCACT |
| Acta2 | CTGACAGAGGCACCACTGAA | CAGAGGCATAGAGGGACAGC |
| Smad4 | AGAGTCTAACGCCACCAGC | TGAAGCTATCTGCAACAGTCCT |
